# Modeling *cis*-regulatory variation in human brain enhancers across a large Parkinson’s Disease cohort

**DOI:** 10.64898/2026.03.15.711881

**Authors:** Olga M. Sigalova, Alexandra Pančíková, Julie De Man, Koen Theunis, Gert J. Hulselmans, Vasileios Konstantakos, Bram Stuyven, Anton De Brabandere, Jarne Geurts, Antonina Mikorska, Shinjini Mukherjee, Sara Abouelasrar Salama, Katy Vandereyken, Kristofer Davie, Lukas Mahieu, Charles H. Adler, Thomas G. Beach, Geidy E. Serrano, Thierry Voet, Jonas Demeulemeester, Stein Aerts

**Author notes:** shared corresponding authors; correspondence. equal contribution (Olga M. Sigalova, Alexandra Pančíková, Julie De Man, Koen Theunis).

## Abstract

Genome-wide association studies (GWAS) have linked more than a hundred non-coding genomic loci to Parkinson’s disease (PD) risk. Deciphering their functional impact on gene regulation requires cell type-aware modeling approaches to assess the effects of sequence variation on enhancer function and target gene expression. To address this challenge, we generated a comprehensive matched dataset from 190 human donors (115 controls and 75 PD), comprising long-read whole-genome sequencing alongside single nucleus multiome atlases (snATAC-seq and snRNA-seq for 3.1 and 1.1 million nuclei respectively) of the anterior cingulate cortex and substantia nigra. By integrating chromatin accessibility quantitative trait loci (caQTL), DNA methylation QTL (meQTL), and allele-specific chromatin accessibility (ASCA), we identified 53,841 high-confidence *cis*-acting genetic variants that modulate cell type-specific enhancer accessibility in one or both brain regions. We further demonstrate that sequence-to-function models can accurately predict the impact of these variants directly from the genomic sequence. Novel explainability approaches allowed stratifying these variants according to their regulatory function, with the majority disrupting specific transcription factor binding sites in a cell type specific manner. Integrating these “enhancer variants” (EV) with eQTL mapping and gene locus modeling linked a subset of EVs to their target genes. Finally, we applied these models to prioritize regulatory variants at known PD GWAS loci, bypassing statistical limitations in rare disease-relevant populations like dopaminergic neurons. All together, we establish a unique resource and new sequence modeling strategies to interpret functional non-coding variation in the human brain.

## Introduction

Human genetic variation is a primary driver of phenotypic diversity and disease susceptibility. While Genome-Wide Association Studies (GWAS) have successfully mapped thousands of loci to complex traits, the vast majority of these variants are non-protein-coding, making their functional interpretation difficult. The primary tasks in tackling this problem are to identify and understand (1) which sequence variants modulate enhancer or promoter function; and (2) which of these variants affect target gene expression, possibly through combinations of distal interactions. Furthermore, as many diseases affect not all cell types, but only one or several, it is expected that many functional cis-regulatory variants have cell type-specific effects, thus requiring assays that work at single-cell or cell type resolution ^1,2^.

Sequence-to-function (S2F) deep learning models have emerged as a powerful approach to decode regulatory logic by predicting chromatin accessibility and gene expression directly from DNA sequence^3–6^. Modeling human genetic variation offers an interesting direction to benchmark and optimize these S2F models. If a model can accurately predict the impact of an unseen variant on regulatory activity, it may capture the underlying regulatory rules. However, systematic validation of these predictions and ground truth datasets, particularly within complex tissues such as the brain, remain limited ^6,7^, and the limitations of such approaches on rare cell types are unclear. Resolving the genomic cis-regulatory code is particularly crucial in the human brain, which exhibits the highest diversity of cell types in the body. This diversity is achieved largely through the action of enhancers and their cognate transcription factor (TF) combination, regulating gene expression in a cell type–specific manner^8,9^.

The difficulty of interpreting non-coding variation is exemplified in Parkinson’s Disease (PD). PD GWAS have identified 134 risk loci that collectively explain 16-36% of the heritable disease risk ^10,11^. While protein-coding mutations in genes such as *SNCA*, *PRKN*, and *PARK7* explain familial cases^12^, few of the GWAS-linked loci have been mechanistically explained^13–17^ because 93% of all lead variants associated to PD-risk are located in the non-coding genome. Identifying the causal variants, which are often not the lead SNP, requires mechanistic interpretations of how they change cell state.

Here, we tackle this problem by generating matching datasets for 115 control and 75 PD donors, of their phased diploid genomes, generated by long-read whole genome sequencing, and their single-cell multiome atlases (snATAC-seq and snRNA-seq) of two human brain regions, the anterior cingulate cortex and the substantia nigra. We then use local S2F models to predict enhancer variants (EV) and validate these using chromatin accessibility QTL (caQTL) and allele-specific chromatin accessibility (ASCA). Subsequently, we link EVs to target genes through expression QTL (eQTL) analyses and assess the performance of large S2F models to predict the effect of EV on gene expression. Finally, we investigate all PD-linked GWAS loci for cis-regulatory variants.

## Results

### Long-read whole genome sequencing on a cohort of donors

To study regulatory variation in the brain, we assembled a cohort of 190 donors and collected tissue from two brain regions: cingulate cortex (CC), a brain region affected in later stages of the disease, and substantia nigra (SN), a brain region first affected in PD^18^ (Fig. 1A, Supplementary Table 1). Cases included 115 neurotypical controls and 75 patients with Parkinson’s disease (PD), obtained from three biobanks (see Methods). Of these donors, 70 were female and 120 were male, and their age at death ranged from 33 to 100 years old, with a median of 75 (median of 80 for PD and 69 for controls; Figure 1B, Supplementary Figure 1A-B). Both age and postmortem interval varied for each biobank, with PMI ranging from 1.83 to 168 hours with a median of 48.6 hours (Supplementary Figure 1C-1D).

**Figure 1:**
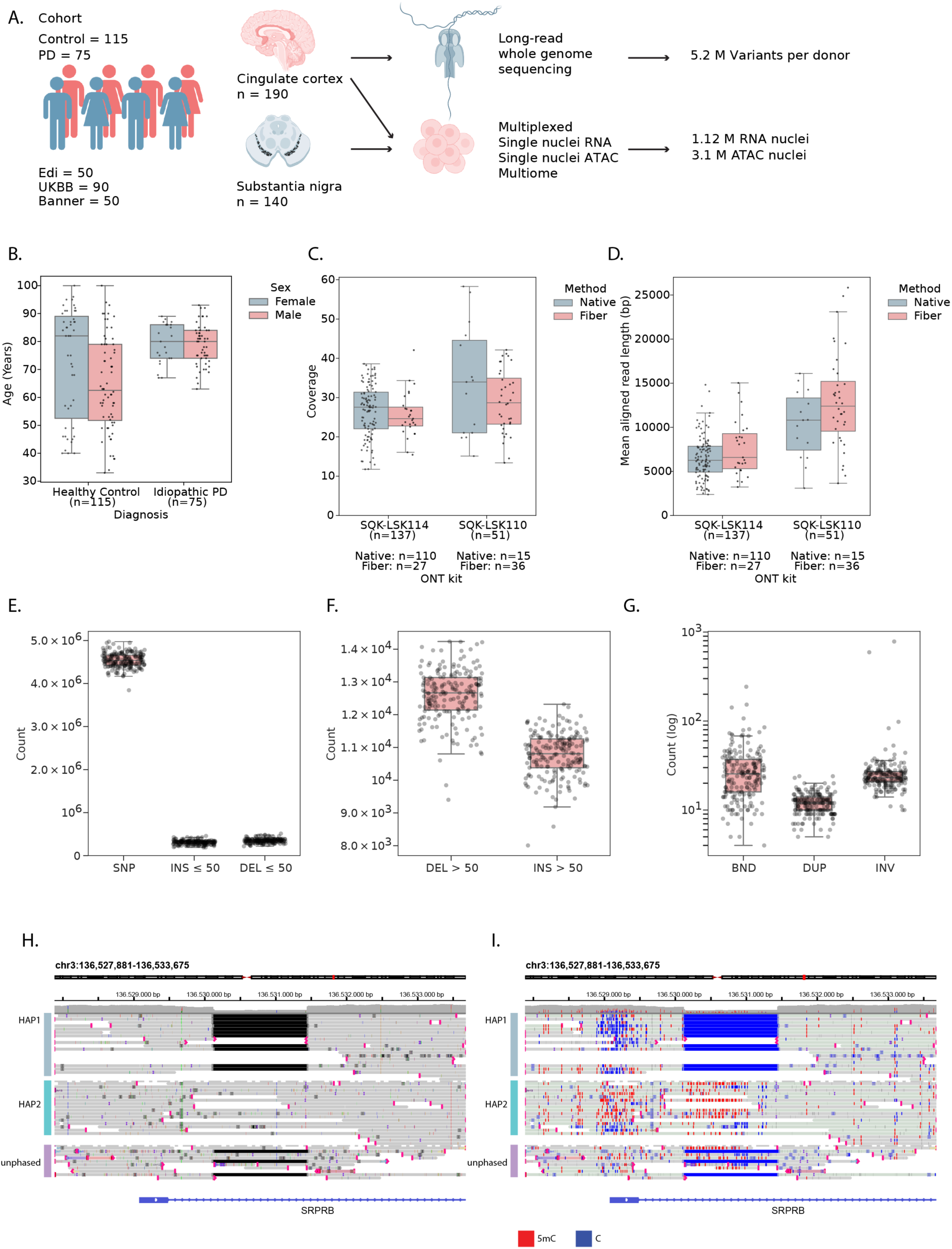
Long-read whole-genome sequencing of 180 post-mortem brains. **A**. Overview of donor cohort and experimental set-up. **B**. Donor age distribution stratified by PD status and sex. **C,D**. Whole-genome coverage (C) and mean aligned read length (D) stratified by sequencing chemistry and method. **E**. Small variant counts per donor. **F**. Variant count for deletions (DEL) and insertions (INS) > 50bps per donor. **G**. Variant counts for breakends, duplications and inversions per donor. **H,I.** Example IGV visualization of phased reads at chr3:136,527,881-136,533,675 of donor ASA_190B, containing 1.3 kb heterozygous deletion (H) and the same view with read CpG methylation status color-encoded showing impact of deletion on methylation status of proximal promoter (red is methylated, blue unmethylated).

We generated long-read whole-genome data from all these cases using Oxford Nanopore Technologies (ONT) nanopore sequencing. This overcomes shortcomings of conventional short-read sequencing, such as missing structural variation and phasing information. We sequenced both native DNA (n = 125), allowing to detect endogenous cytosine (hydroxy)methylation, and DNA from nuclei treated with the N6-adenosine methyltransferase Hia5 (Fiber-Seq^19^) to additionally reveal base pair-resolution open chromatin information via the modified base read output (n = 63) (Methods). WGS data was processed with the in-house developed Nextflow pipeline NanoWGS (Supplementary figure 1G, Supplementary Table 2, Methods). We obtained an average sequencing coverage of 27.5X with an aligned read length of 8.2 kb (Figure 1C-1D). We ascertained the ancestry of donors: 185 donors were of European ancestry, 2 of admixed American ancestry and 1 of East Asian ancestry (Supplementary Figure 1H), and we benchmarked the accuracy of SNP and indel calling of different ONT chemistries (Supplementary Figure 1E-F, Methods). After alignment to the T2T-CHM13v2.0 reference genome, we detected an average of 4.54M single nucleotide polymorphisms (SNPs), ∼657k indels ≤ 50 bps (Figure 1E, INS: 309,129, DEL: 347,873) and ∼23K structural variants (SVs; variants of size > 50 bps, 12,584 deletions, 10,769 insertions, 32 inversions (INV), 12 duplications (DUP), 31 unannotated breakends (BND) per genome, in line with recent literature^20^ (Figure 1F-1I). We observed peaks in variant numbers around 300 bps, 2-3 kbps and 6-7kbps, representing ALU, SVA and LINE-1 elements, respectively (Supplementary Figure 1J-K). Taken together, we were able to generate high quality multi-modal long read data for our cohort, resulting in phased diploid genomes and confident SNP, indel, and SV calls.

### Matched single-nucleus ATAC and RNA atlases of human substantia nigra and cingulate cortex

We performed single-nucleus ATAC and RNA sequencing of the same donors, for the CC (115 controls, 75 PD patients) and SN (90 controls, 50 PD) (Figure 1A, Methods), using a combination of sn-multiome (snRNA+snATAC), snRNA-seq, and snATAC-seq (see Methods, Supplementary Figure 2S & 2T). Cell type annotation was performed with scANVI models trained to transfer labels from external datasets^21^, combined with manual curation (Figure 2A and 2C). For the SN, we obtained 376,394 RNA nuclei with a mean of 1,969 genes and 4,696 UMIs per nucleus (Supplementary Figure 2D-F & Supplementary Figure 2J-L) and identified the expected non-neuronal cell types (Fig 2E, Supplementary Figure 2W). Neuron types were annotated as dopaminergic neurons (DopaN), gabaergic neurons (GabaN) and glutamatergic neurons (GlutaN, Figure 2E). Only 18.2% of DopaN derive from PD donors as they are gradually lost during the disease, while the number of GabaN and GlutaN is comparable between PD and control donors. In the CC, we obtained 751,541 high quality nuclei with a mean of 2,541 genes and 7,150 UMIs per nucleus (Supplementary Figure 2A-C & Supplementary Figure 2G-I). Here we observe the expected cortical cell types including all the layered excitatory neurons and all inhibitory neuron types (Supplementary Figure 2U, Supplementary Figure 3A).

**Figure 2:**
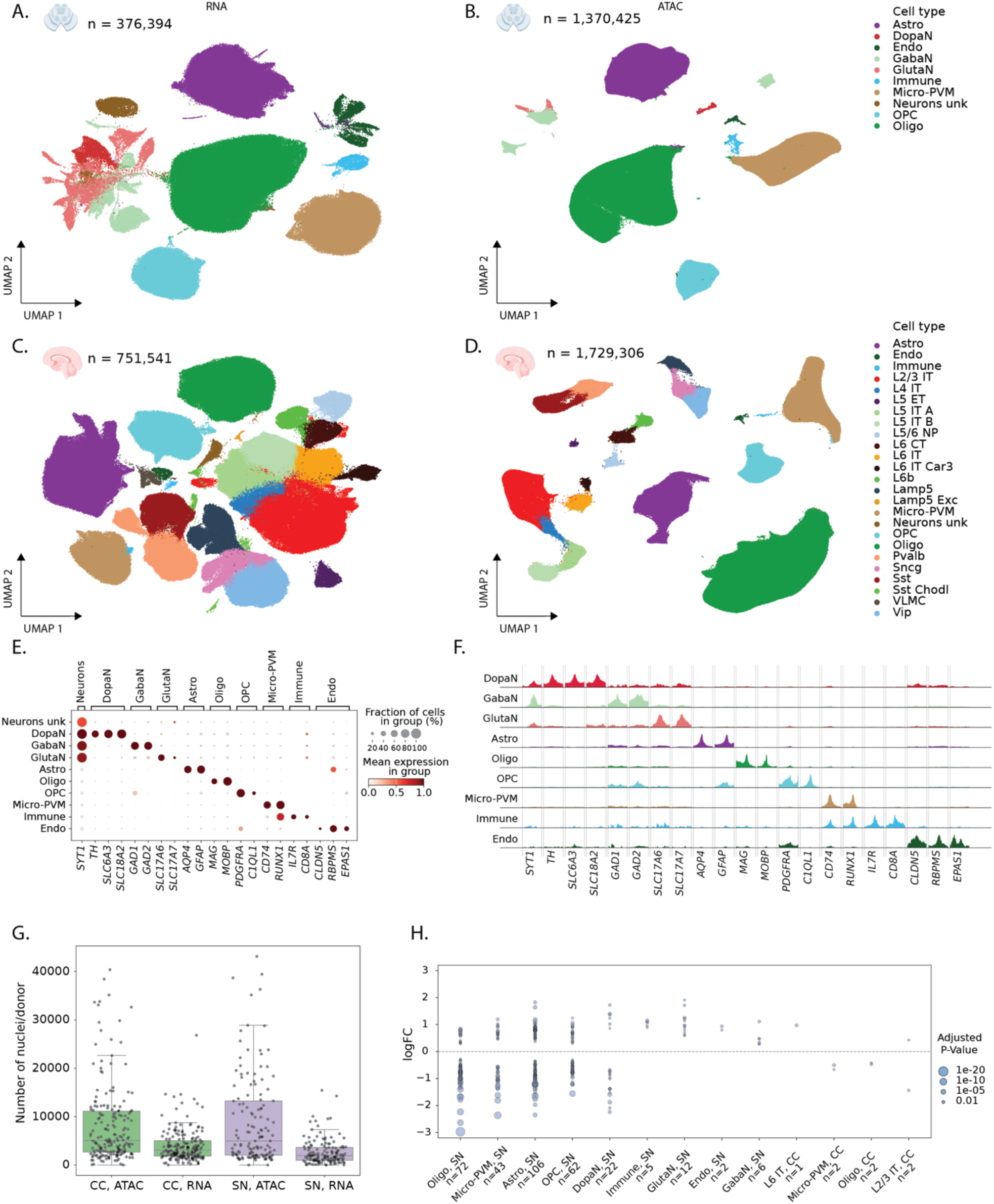
**Single-cell multiomic atlases of substantia nigra and cingulate cortex. A,B**. UMAPs of substantia nigra single nuclei gene expression (A) and chromatin accessibility (B) colored by cell type. **C,D**. UMAPs of cingulate cortex single nuclei gene expression (C) and chromatin accessibility (D) colored by cell type. **E**. Average gene expression of select marker genes in substantia nigra cell types. **F**. Pseudobulk chromatin accessibility profiles in proximity of select marker genes in substantia nigra cell types. **G**. Number of nuclei per donor in each brain region and modality. **H**. Log2 fold change of all differentially expressed genes reported per cell type (p-adj < 0.05).

To process the snATAC-seq fragments, we performed quality control, topic modeling and clustering with pycisTopic ^22^. We annotated nuclei using matched multiome RNA annotations and extrapolated annotations to the cluster level using a majority voting strategy (see Methods, Figure 2B, Figure 2D, Supplementary Figure 2U-W). For CC, this resulted in an atlas of 1,729,306 high-quality, annotated nuclei, with 1,707,448 high confidence chromatin accessibility peaks (Supplementary Figure 2M-O,T, Supplementary Figure 3B). For the SN, the atlas contains 1,370,425 high-quality, annotated nuclei and 1,292,285 chromatin accessibility peaks (Figure 2F, Supplementary Figure 2P-R,T). Calling differentially accessible regions per cell type identified 380,770 and 115,474 cell-type specific regions for CC and SN, respectively.

To decrease sequencing costs and batch effects, we pooled 2-10 donors in each experiment. We then used donor-specific variants to demultiplex cells per donor (matching 98.4% between ATAC and RNA for the multiome runs, Methods). This yielded an average of 3,464 nuclei per donor in snRNA (Figure 2G, range 4-26,845) and 8,547 nuclei per donor in snATAC (Figure 2G, range 1-43,410). Next, we performed cell type-specific differential gene expression between healthy and PD donors using dreamlet^23^. After multiple testing correction (Benjamini-Hochberg adjusted p-value < 0.05), we detect 337 differentially expressed genes across 13 cell types (Figure 2H). 330 of these differentially expressed genes are in SN cell types, while only 7 genes are reported to be differentially expressed in CC (Supplementary Table 3). While CC is affected in later stages of PD^18^, our analysis did not yield many differentially expressed genes in this region.

### Cis-acting genetic variants affect cell type specific chromatin accessibility

To comprehensively quantify the impact of genetic variation on chromatin accessibility, we applied two complementary approaches: chromatin accessibility quantitative trait loci (caQTL) mapping and allele-specific chromatin accessibility (ASCA) analysis (Figure 3A-B, Methods). caQTL mapping tests for associations between genotype and chromatin accessibility across donors, capturing population-level effects, whereas ASCA measures allelic imbalance in chromatin accessibility within heterozygous individuals, providing an internal, within-sample control that helps account for trans-acting variation and technical variability^24,25^. Together, these approaches provide orthogonal and mutually reinforcing evidence, with ASCA adding an additional layer of control to support interpretation of caQTL signals.

**Figure 3.**
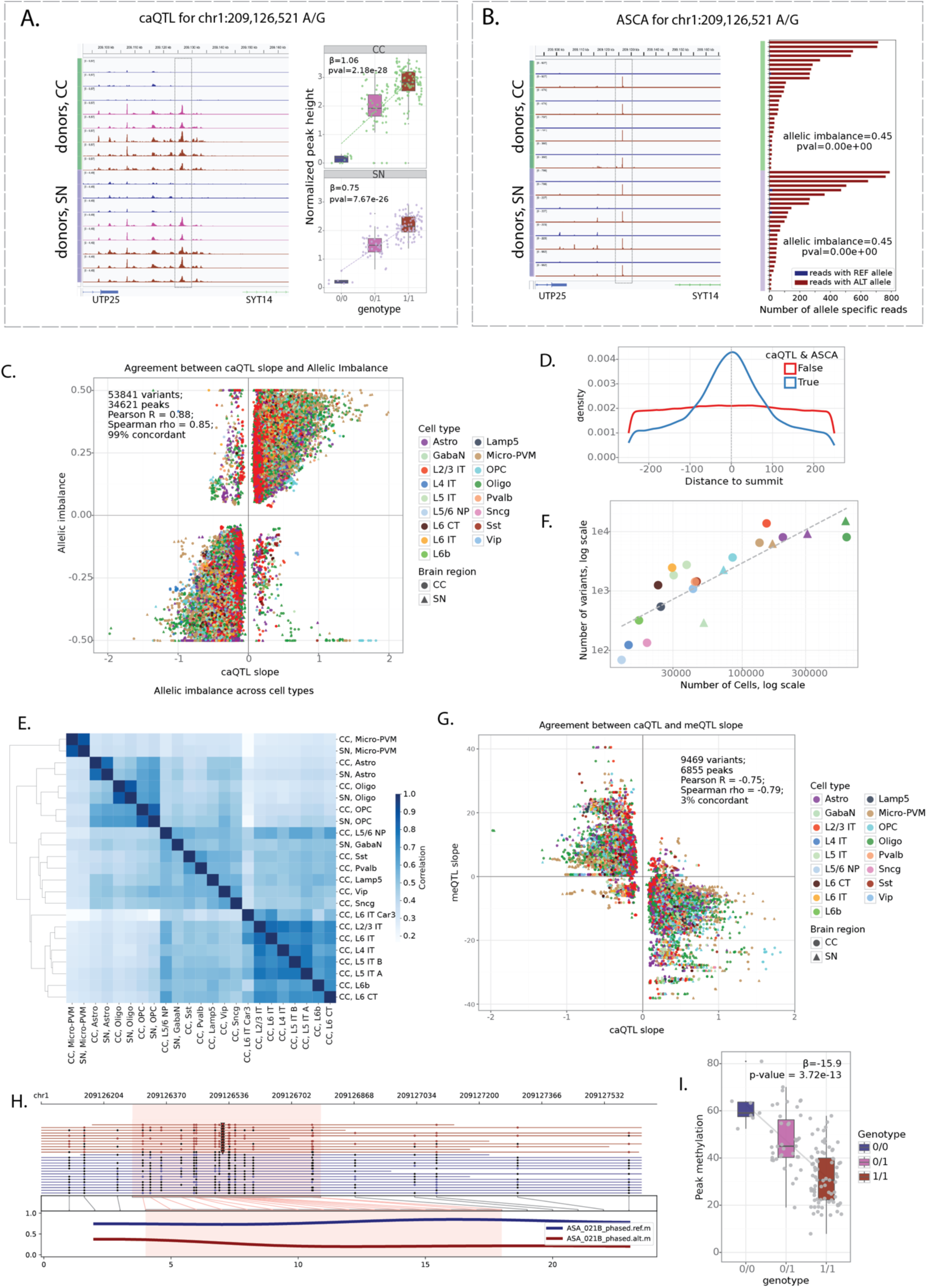
caQTL and ASCA profiling reveals cis-acting cell type-specific chromatin accessibility variants. **A**. Chromatin accessibility upstream of *SYT14* in Oligodendrocytes across donors with different genotypes for SNP chr1:209,126,521 (A/G). Browser tracks are shown for eight donors in CC and SN. The corresponding ATAC peak is boxed. Boxplots show normalized peak height for all quantified donors, matched track and boxplot colors correspond to donor genotype. **B.** ASCA in Oligodendrocytes of the same genomic locus as in A. Allele-specific tracks are shown for four heterozygous donors in CC and SN (two tracks per donor, only reads overlapping heterozygous variants are considered. The corresponding ATAC peak is boxed. Barplots show the number of reads from REF and ALT alleles for all heterozygous donors with ≥10 phased reads for the corresponding peak in Oligodendrocytes. **C**. Concordance between caQTL slope and allelic imbalance for variants significant for both caQTL and ASCA tests at FDR<0.1. If a variant is significant in multiple cell types (independent), it is represented by multiple dots. **D**. Distribution of variants’ locations relative to peak summit. The variant with the smallest caQTL *p*-value per peak is selected. All quantified peaks in all cell type are considered (1.5M peaks total). Peaks with caQTL & ASCA variants are shown in blue. **E**. Correlation of allelic imbalances across cell types. All caQTL & ASCA variants in at least one cell type are considered, allelic imbalances are compared across all cell types where a variant is quantified (regardless of significance). **F**. Number of caQTL & ASCA variants vs. total number of cells per cell type (across all donors). **G**. Concordance between caQTL and mQTL slopes for caQTL & ASCA variants with mQTL FDR<0.1. **H**. Methylartist visualization of donor ASA_021 phased ONT WGS reads at the *SYT14* SNP and mQTL from A-B (chr1:209,126,521 (A/G)). The variant is depicted as a triangle and the allele-specifically methylated region highlighted. Methylated CpGs are full black circles, while unmethylated ones are empty. **I**. Boxplot with mean CpG methylation levels in region chr1:209126280-209126780 stratified by donor genotype at the same *SYT14* variant.

caQTLs were identified using tensorQTL^26^ incorporating sex, age, donor biobank, sequencing modality, five genotype principal components, and thirty data principal components as covariates. ASCA analysis was performed using the WASP framework^27^, extended to include indels and adapted to account for cell barcodes and phasing-block information (Methods). This pipeline includes mappability-bias correction and leverages the allele-specific component of the Combined Haplotype Test (AS-CHT) to obtain robust estimates of allelic imbalance. We restricted our analysis to SNPs and indels located within the regions of accessible chromatin (consensus peaks) of CC and SN, yielding 2.2 million variants (2,105,788 SNPs and 128,503 indels) tested by both caQTL and ASCA in at least one cell type (Methods).

Figure 3A-B shows an example SNP at chr1:209,126,521 (A/G) associated with differentially accessible chromatin by both caQTL and ASCA analyses. The corresponding region is highly accessible in oligodendrocytes and lies upstream of the *SYT14* gene (Supplementary Figure 4A). Donors homozygous for the reference allele (A/A) have lower accessibility in the region compared to heterozygous (A/G) donors and those homozygous for the alternative allele (G/G) (Figure 3A). Furthermore, in heterozygous donors, most reads come from the haplotype containing the alternative allele (Figure 3B), supporting the *cis*-effect of the genetic variant.

caQTL and ASCA analyses gave highly concordant estimates of variant effects, even at relaxed FDR thresholds (Supplementary Figure 4B-C). We identified 53,841 variants within 34,621 consensus peaks with FDR < 0.1 by both caQTL and ASCA analyses (referred to as *caQTL-ASCA* variants), including 50,860 SNPs and 2,981 indels (Supplementary Table 4). caQTL slopes and allelic imbalance measurements were strongly correlated (Pearson’s *r* = 0.88), and 99% of variants exhibited the same direction of effect (Figure 3C). Significant variants had strong enrichment towards snATAC peak summits (Figure 3D), suggesting that these variants are more likely to directly disrupt TF binding sites. Notably, variant quantification was based on all reads within peaks (Methods), indicating that this enrichment is unlikely to be explained by read-coverage biases around peak summits. Furthermore, variant effects were correlated across related cell types (e.g. L2/3-L6 neurons in CC) and between the same cell types across brain regions (e.g. microglia in CC and SN), even though all quantifications and testing were done independently for each cell type (Figure 3E, Supplementary figure 4D). Thus, our combined approach identifies high confidence reproducible set of cis-acting genetic variants.

The number of significant variants scaled with the number of cells per cell type, indicating that statistical power to detect genetic effects is limited in rare cell types (Figure 3F), consistent with previous single-cell QTL studies^28–32^. Consequently, most caQTL-ASCA variants were identified in common cell types, whereas some rare cell types (including dopaminergic and glutamatergic neurons in SN) were excluded from this analysis because of the limited number of cells per donor. To approximate genetic effects in these populations, we additionally performed caQTL analysis in broader neuronal classes (excitatory and inhibitory neurons in CC, and all neurons in SN). This resulted in identifying 41,686 variants with FDR<0.1 in broad neuronal classes, with caQTL slopes highly correlated with the underlying granular cell types (Supplementary Figure 4E).

We next analyzed CpG methylation using bulk ONT WGS. As an initial quality control of the methylation calls, we created CpG methylation pileups for samples with WGS data derived from CC, we compared the methylation levels at TSS of highly and lowly expressed genes and confirmed clear hypomethylation at highly expressed genes and high methylation at lowly expressed genes in all samples (Supplementary Figure 5A). Of note, we observe differences in CpG methylation levels between the ONT r9 and r10 flow cells and between Fiber-seq samples and samples without exogenous labelling. Looking at the donor-specific pileups of all reads around TSS and CTCFs, we see clear hypomethylation which is more pronounced downstream of TSS at highly expressed genes (Supplementary Figure 5B) and decrease in accessibility with clear nucleosome patterning around CTCF motifs (Supplementary Figure 5C). We also observe clear haplotype-specific methylation at imprinted regions (Supplementary Figure 5D).

We performed bulk meQTL mapping for 125 donors with tensorQTL^26^ (Methods) and identified 57,951 unique regions where methylation was significantly affected by genetic variants present in our cohort (FDR<0.05). Out of 53,841 caQTL & ASCA variants, 9,469 (17.6%) variants also showed effect on methylation (meQTL nominal p-value < 1e-3), with a strong negative relationship between meQTL and caQTL signals (Pearson’s *r* =-0.78, 96% discordant, Figure 3G), indicating reduced CpG methylation on the more accessible haplotypes and vice versa (Supplementary Table 5). For example, allele G of the SNP at chr1:209,126,521 associated with increased chromatin accessibility in oligodendrocytes (Figure 3A-B) was also found to be associated with decreased methylation, detected in bulk (Figure 3H-I).

### Sequence-to-function models predict variant effects on chromatin accessibility

Our analysis identified a consistent set of genetic variants affecting chromatin accessibility and DNA methylation in the donor cohort. Next, we asked whether these effects could be predicted directly from the genome sequence using sequence-to-function models, without requiring prior information about genetic variation across individuals. To address this question, we trained CREsted^4^ peak regression models, which take as input the reference genome sequences and as output predict cell type-specific chromatin accessibility. The models were trained separately on consensus peaks of CC and SN (DeepCC and DeepSN models) and subsequently finetuned on highly variable regions (Methods). The trained models showed good performance for predicting cell type-specific chromatin accessibility, with Spearman correlation of 0.76 in CC and 0.82 in SN between predictions and mean ATAC signal on the test set of highly variable regions across cell types (Supplementary Figure 6A-F). We then used the trained models to predict the regulatory effects of ∼7M SNPs and ∼800K indels from the donor cohort using *in silico* mutagenesis (ISM), i.e. by introducing the alternative allele of each variant into the reference sequence of the corresponding peak and calculating the log-fold change between alternative and reference prediction in all cell types.

We found strong concordance between variant effects predicted by CREsted models and those measured in donors by ASCA and caQTL (Figure 4A-C, Supplementary Figure 6G, Supplementary Table 4). Among 34,621 caQTL-ASCA peaks, 83% showed concordant effect directions between the CREsted prediction (variant logFC) and the observed allelic imbalance in donors (Spearman’s rho = 0.71), with concordance increasing to 99% (Spearman’s rho = 0.81) for the peaks with stronger predicted effects according to the deep learning models (absolute logFC > 0.5). Importantly, both models were trained on the reference human genome and thus do not incorporate information about genetic variation present in the donor cohort. This indicates that variant effect predictions are based on general biology learned by the models and not overfit to donor-specific data. To illustrate this, we also used the previously published DeepHumanBrain model^33^ trained on external human brain data^34^ and scored variants in the cell types matching between the two datasets. Variant scores were concordant between the models, with Spearman’s rho of 0.81 on the 34,141 caQTL-ASCA variants with absolute logFC above 0.05 by both models (Figure 4D). Furthermore, high agreement (Spearman’s rho = 0.92 on 18,221 variants with matching cell types and absolute CREsted logFC above 0.05) was observed between DeepCC and the DeepBICCN2 model^4^ trained on mouse cortex data^35^, suggesting conserved and sequence explainable features underlying the models’ predictions (Supplementary Figure 6H).

**Figure 4.**
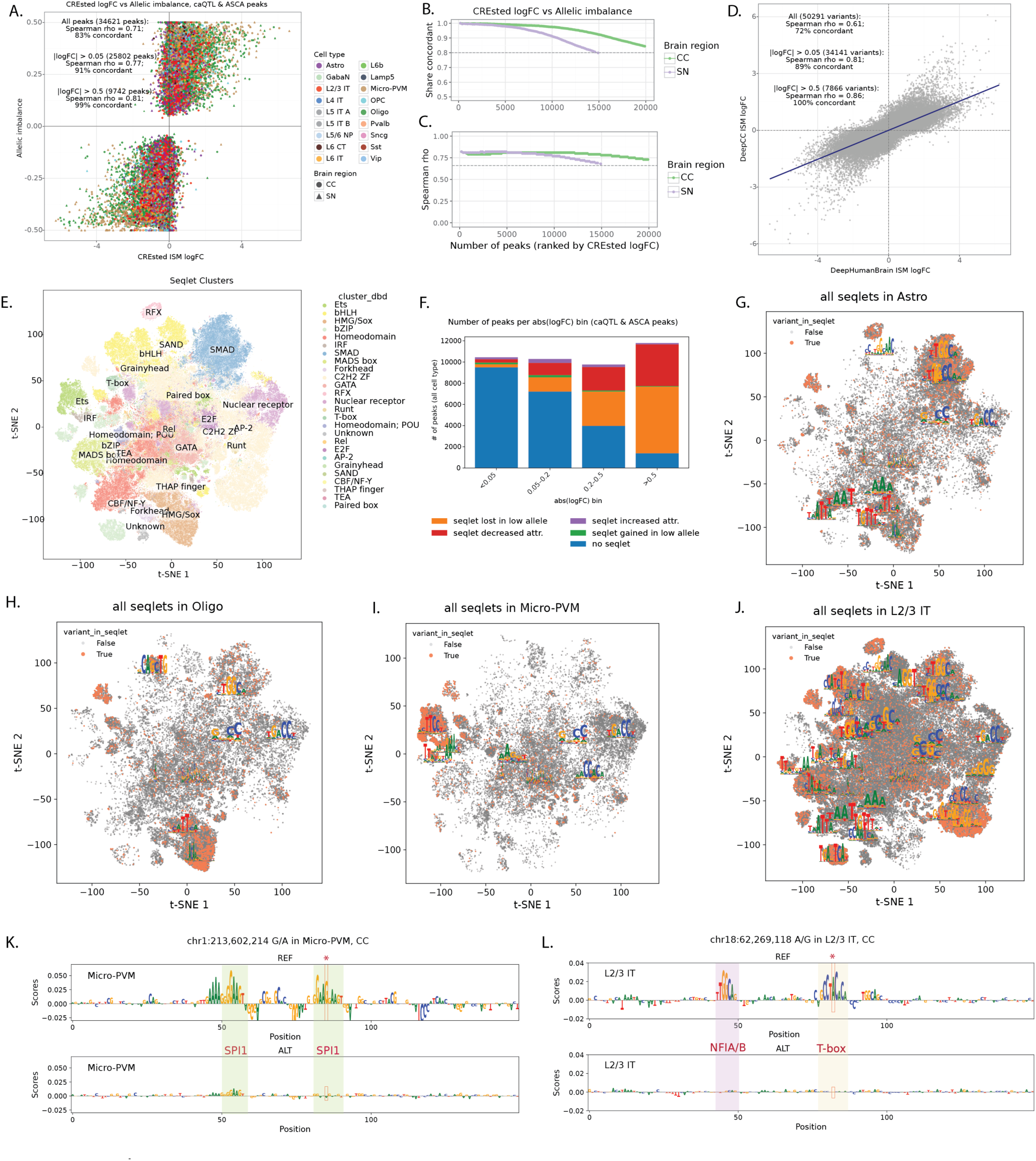
Modeling cis-regulatory variation in the brain. **A.** Agreement between CREsted *in-situ* mutagenesis (ISM) logFC and allelic imbalance for all 34,621 caQTL-ASCA peaks. **B.** Share of peaks with concordant (same sign) CREsted logFC and allelic imbalance for different number of top CREsted predicted peaks (x axis). **C.** Spearman correlation between CREsted logFC and allelic imbalance for different number of top CREsted predicted peaks (x axis). **D.** Agreement between DeepHumanBrain logFC vs DeepCC logFC for variants tested. **E.** tSNE dimensionality reduction of 663,034 seqlets extracted from caQTL-ASCA peaks in all cell types (high and low allele) colored by TF-family annotation. **F.** caQTL-ASCA peaks binned by CREsted logFC and categorized by variant effects on seqlets (all cell types combined). **G-J.** tSNE dimensionality reduction of all seqlets in Astrocytes (G), Oligodendrocytes (H), Micro-PVM (I), and L2/3 IT neurons (J), colored by whether the affected variant was present in the seqlet with position weight matrix logos for clusters with at least 1000 seqlets for a given cell type. **K,L.** DeepExplainer of REF (top) and ALT (bottom) allele for variant chr1:213,602,214 G/A in Micro-PVM, CC (K) and variant chr18:62269118 A/G in L2/3 IT, CC. Tested variants are highlighted by a red box and star. The affected motifs are highlighted and annotated with their TF-family.

Since *cis*-acting non-coding genetic variation is expected to act mostly through the disruption of TF binding sites^36,37^, we next analyzed sequence motifs affected by the caQTL & ASCA variants according to CREsted models using TF-MINDI package^38^ (Methods). For this analysis we extracted per-nucleotide attribution scores for reference and alternative peak sequences and used them to call seqlets^39^ (short stretches of nucleotides with high attribution, these represent candidate TF binding sites). These seqlets were then clustered based on sequence similarity and mapped to a database of known TF binding motifs (Figure 4E). We observed an overall decreased number of seqlets called in the allele with lower CREsted predictions (‘low allele’) (384,194 in the high allele, 278,840 in the low allele). Overall, for about 50% of caQTL-ASCA peaks, the top CREsted variants are predicted to directly affect TF motifs leading either to seqlet loss or decreased attribution in the low allele (Figure 4F). In particular, about 90% of peaks with strong CREsted predictions (abs. logFC > 0.5) had variants directly affecting seqlets. Variants with weak CREsted predictions (abs. logFC < 0.2) mostly couldn’t be linked directly to affected TF motifs, even though model predictions were still generally concordant with observed effects in donors (Figure 4A-C). Those variants could potentially act on low affinity binding sites, flanking motif sequences or other sequence features influencing general chromatin accessibility which could be captured by the sequence-to-function models but don’t directly map to canonical TF motifs. Seqlets directly affected by caQTL-ASCA variants showed strong cell type-specific patterns (Figure 4G-J, Supplementary Figure 7A) and mapped to known cell type-specific regulators including SPI1 motifs (Ets family) for microglia and SOX family motifs for oligodendrocytes.

Figures 4K-L show examples of two variants with concordant CREsted predictions and observed effects on chromatin accessibility in donors. SNP at chr1:213,602,214 falls into region specifically accessible in Microglia, and alternative allele (A) is predicted to reduce chromatin accessibility in that cell type in both CC and SN due to disruption of SPI1 binding motif (Figure 4K), which agrees with the observed effects on chromatin accessibility by both caQTL and ASCA (Supplementary Figure 7B). SNP at chr18:62,269,118 is predicted to affect region with broader accessibility in different subtypes of excitatory neurons in CC due to disruption of T-box motif (Figure 4L). Concordantly, this variant has significant effect observed in donors according to both caQTL and ASCA in L2/3 IT, L5 IT B, and L6 IT neurons (Supplementary Figure 7C). In L5 IT A, L5 ET, L6 CT, and L6 IT Car3 neurons observed caQTL slopes are concordant with CREsted predictions, but they don’t pass stringent caQTL & ASCA significance criteria. Thus, deep learning predictions can potentially increase power of the standard statistical tests, especially in the rare cell types, which is supported by the observation that the number of CREsted variants with strong effects sizes didn’t scale with the number of cells per cell type (Figure 5A), as opposed to caQTL and ASCA (Figure 3F).

**Figure 5.**
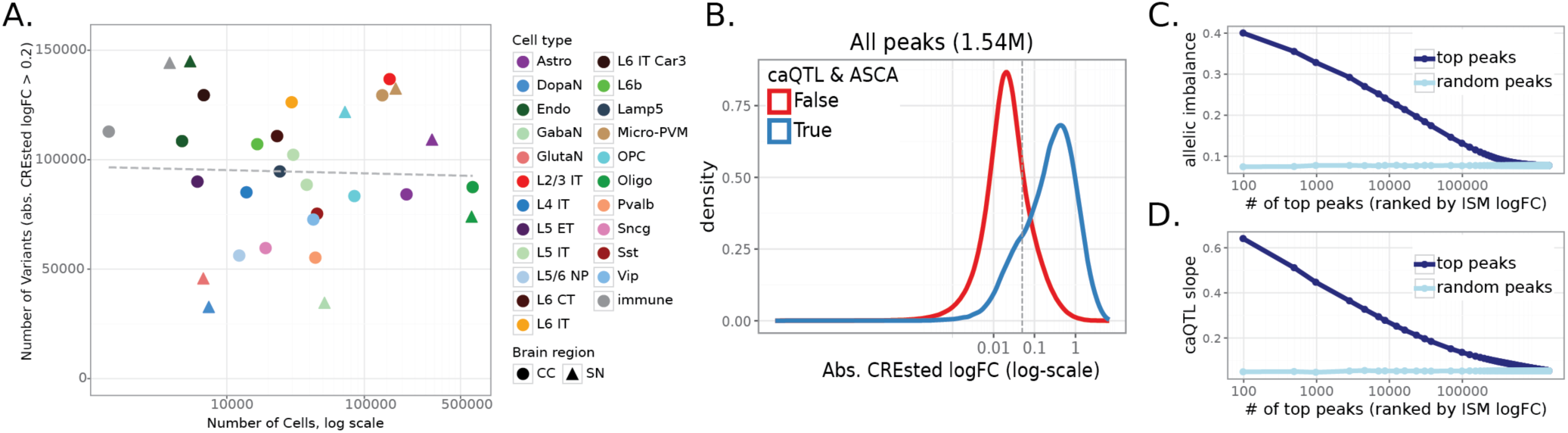
CREsted models increase power of variant discovery. **A.** Number of cells per cell type *vs.* the number of variants with absolute crested prediction logFC > 0.2. **B.** CREsted absolute logFC for all peaks stratified by whether or not the peak contained the caQTL-ASCA variant. **C-D.** Rank plot of average allelic imbalances (C) and caQTL slopes (D) for top peaks ranked by CREsted logFC (dark blue) vs. same number of randomly selected peaks (light blue).

We next asked whether CREsted predictions can recover caQTL & ASCA variants directly from the genome sequence. Across all 2.2 million variants quantified in at least one cell type, caQTL & ASCA variants exhibited higher CREsted scores than variants without detectable donor effects, with the two distributions clearly separated by the CREsted logFC threshold of about 0.1 (Figure 5B). Furthermore, variants with higher ranks by CREsted ISM had higher absolute allelic imbalance (Figure 5C) and caQTL slope (Figure 5D) compared to the lower scoring variants, even when not statistically significant by either of these tests (Supplementary Figure 8A-B). Strong CREsted predicted variants also showed high concordance between CREsted logFC and meQTL slope (Supplementary Figure 8C-D) and caQTL slope in broad neuronal cell types (Supplementary Figure 8E-F) regardless of whether the variant was in the caQTL-ASCA callset or not. Taken together, these results suggest that deep learning models of chromatin accessibility can correctly predict variant effects in donors and thereby can increase power of variant discovery, including cases with low cell and donor counts which do not pass significance thresholds in standard statistical approaches such as caQTL and ASCA.

### Linking and predicting regulatory genetic variation to changes in gene expression

To link the effects of non-coding variation to gene expression, we performed expression quantitative trait locus (eQTL) mapping. Like for caQTL and meQTL analyses, eQTLs were identified using tensorQTL framework. In short, we generated cell type-specific donor pseudobulk expression profiles from the snRNA-seq data and tested all the genes and all the SNP and indels (≤ 50 bp) that were passing the quality filtering within 1 Mb of the gene body. We used the first five genotype principal components, age, sex, biobank and sequencing method as covariates. Of note, since the number of nuclei and donors was limited for individual SN neuronal cell types, to increase the statistical power, we combined these neurons together into a single cell type class for the eQTL testing (Methods). Using this approach, we identified 3,889 unique protein coding genes with a significant eQTL at FDR<0.05 for 22 different cell types in two brain regions (Supplementary Figure 9A). Similar to caQTL results, the detection power scaled with the number of nuclei available per cell type (Supplementary Figure 9B).

Next, we asked to what extent the above identified enhancer variants associate with changes in gene expression. We therefore overlapped the entire caQTL-ASCA callset with eQTL variants with FDR < 0.05 (Supplementary Fable 6). This identified 1,121 genes that have a variant affecting both chromatin accessibility and gene expression (4,717 variants), with Spearman correlation of 0.66 between eQTL and caQTL slope and 86% of variants having concordant direction of effects on these two modalities (Figure 6A). Furthermore, this concordance was higher for a more stringent eQTL set, e.g. Spearman’s rho = 0.73, 92% concordant for 562 genes with FDR < 0.001 (Figure 6B-C).

**Figure 6.**
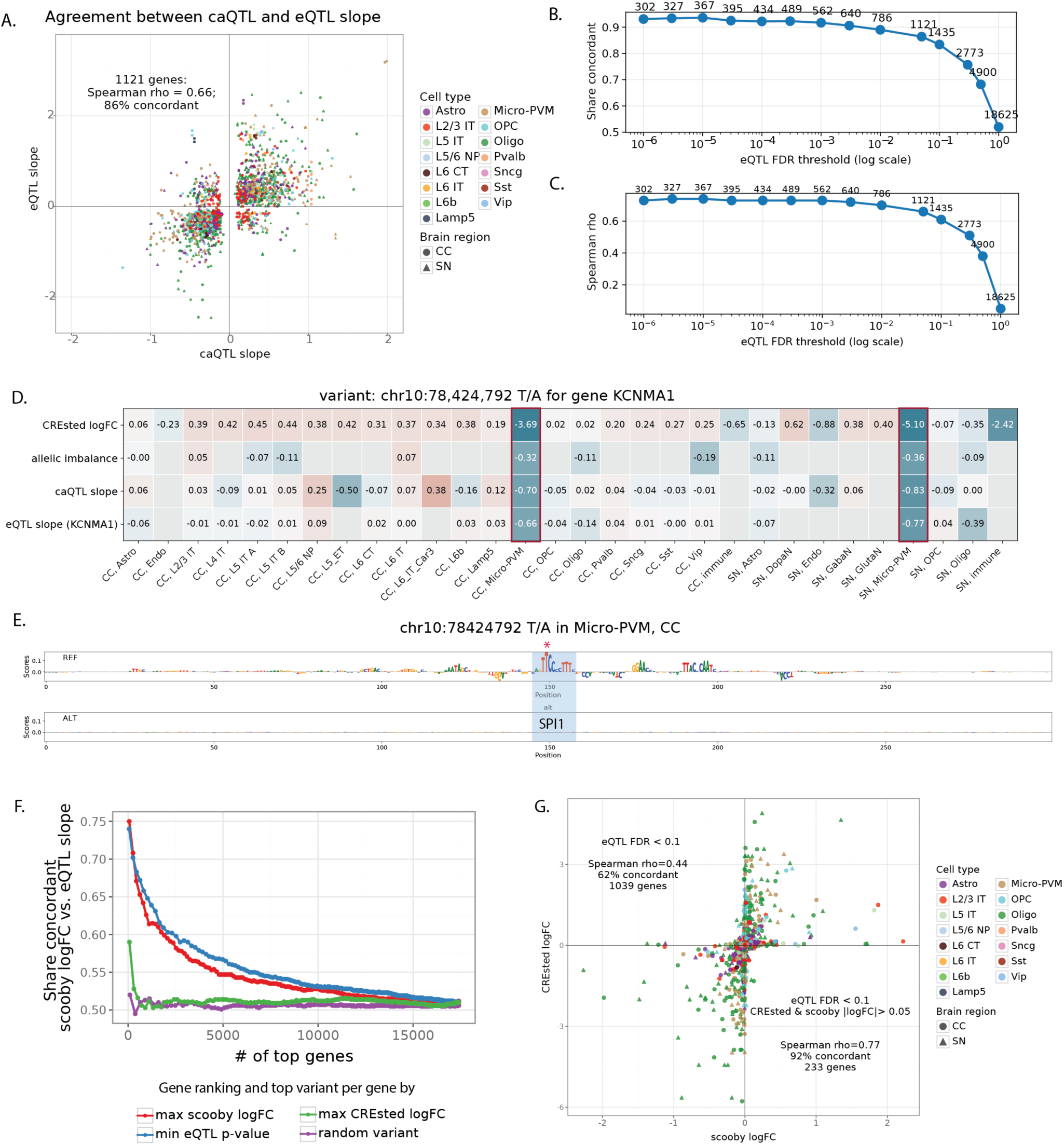
**Linking regulatory variation to gene expression changes**. **A.** Agreement between eQTL and caQTL slope for the overlap of the caQTL-ASCA variant call set with eQTLs with FDR < 0.05. **B**. Share of concordant (eQTL vs. caQTL slope) ASCA-caQTL callset variants overlapping with eQTL variants at different eQTL FDR adjusted p-values. **C**. Spearman correlation between the caQTL and eQTL slopes of ASCA/caQTL call set variants overlapping with eQTL variants thresholded at different FDR-adjusted *p*-values. **D**. Table with Crested logFC, allelic imbalance, caQTL slope, and eQTL slope for *KCNMA1* in all cell types tested for variant chr10:78,424,792 (T/A). Effect sizes in Micro-PVM in SN and CC are highlighted. **E**. DeepExplainer plots for REF and ALT alleles in microglia for chr10:78,424,792 (T/A). Variant location is starred. **F**. Rank plot of share of concordance between scooby logFC and eQTL slope for top genes for all ASCA/caQTL variants. Different lines represent different gene rankings and top variant per gene selection methods. **G**. Agreement between scooby logFC and CREsted logFC for variants with FDR-adjusted eQTL *p*-value < 0.1

Figure 6D-E shows an example of enhancer variant (EV) at chr10:78,424,792 falling in the intron of *KCNMA1* (encoding alpha subunit of the calcium-activated potassium (BK) channel) gene. The SNP is predicted to disrupt SPI1 motif leading to specific loss of chromatin accessibility in microglia. Concordantly, this variant was identified as significant caQTL, ASCA and eQTL of *KCNMA* in the donor cohort, making it a strong candidate EV affecting cell type specific gene expression.

Above, CREsted models could accurately predict local chromatin accessibility changes, and vice versa, local chromatin accessibility changes could be almost entirely explained by sequence changes under the ATAC peak. Now we ask whether larger S2F models that are trained on large input sequences to predict gene expression, such as Borzoi and AlphaGenome^6,40^ can predict the effect of SNPs and indels on gene expression. Our set of 1,121 EV-gene pairs provide an ideal test set to benchmark such methods. To adapt such a model to our data, we opted for a fine-tuning strategy with scooby, based on the Borzoi base model^41^. Particularly, we trained a CC and SN model with Poisson-MultiVI single-cell embeddings^42^ (Supplementary Figure 9C-D). We then used the fine-tuned models to predict the effect of variants on gene expression, for each cell type level (see Methods). We first evaluated the scooby model performance by calculating Pearson correlation between predicted and observed gene expression profiles of all cells within a cell type, for each gene that overlaps the test set sequences (mean Pearson correlation for both models was 0.88, Supplementary figure 9E-F). Correlating predicted and observed log2-transformed gene expression values for each test set gene over all cell types, after mean subtraction across genes and cells, resulted in 0.67 mean Pearson correlation for the CC model, and 0.61 mean Pearson correlation for the SN (Supplementary figure 9G-H).

We used scooby models to score effects of caQTL-ASCA SNPs on all genes where the SNP is within 524 kb window, centered on the gene body (Supplementary Table 7). We found that scooby predictions were generally concordant with eQTL for the variants with strong effects, but that agreement quickly dropped for the less significant SNPs (Figure 6F). CREsted models alone could not predict variant effects on gene expression, which is expected since they were trained on local chromatin accessibility data. However, for the subset of eQTL variants with both scooby and CREsted predictions (233 genes with CREsted and scooby logFC > 0.05), both models showed good concordance (Spearman’s rho = 0.77, 92% concordant) implying similar sequence features underlying variant effects on these two data modalities (Figure 6G).

### Screening PD-associated loci for regulatory variation

We next investigated whether our framework could help to explain disease-relevant loci. Particularly, we screened all loci that were identified in the recent PD GWAS study^11^. Based on linkage disequilibrium, we defined regions around the 157 lead variants (Methods). Of these regions, 43% (*n*=67) contained caQTL-ASCA variants, from which 76% (*n*=51) were also predicted (abs. logFC > 0.2) to affect chromatin accessibility by CREsted models, and 10 blocks contained EVs that were also eQTLs with FDR < 0.05 (Figure 7A-B, Supplementary Table 8). Some of the associated genes are located in well characterized PD loci including *HLA* gene locus^43–45^ and H1/H2 haplotype^46–48^ (locus characterized by a 1Mb inversion spanning 15 genes with strong links to PD risk). Some other genes were previously linked to neurodegenerative diseases, including *NOTUM*^49^ and *SLC2A13*^50^ (Figure 7B, Supplementary Figure 10A-G). However, only two loci contained EVs with direct GWAS evidence (p-value < 5e-8) linked to *TMPRSS5* and five genes in H1/H2 haplotype (Figure 7B, bold).

**Figure 7.**
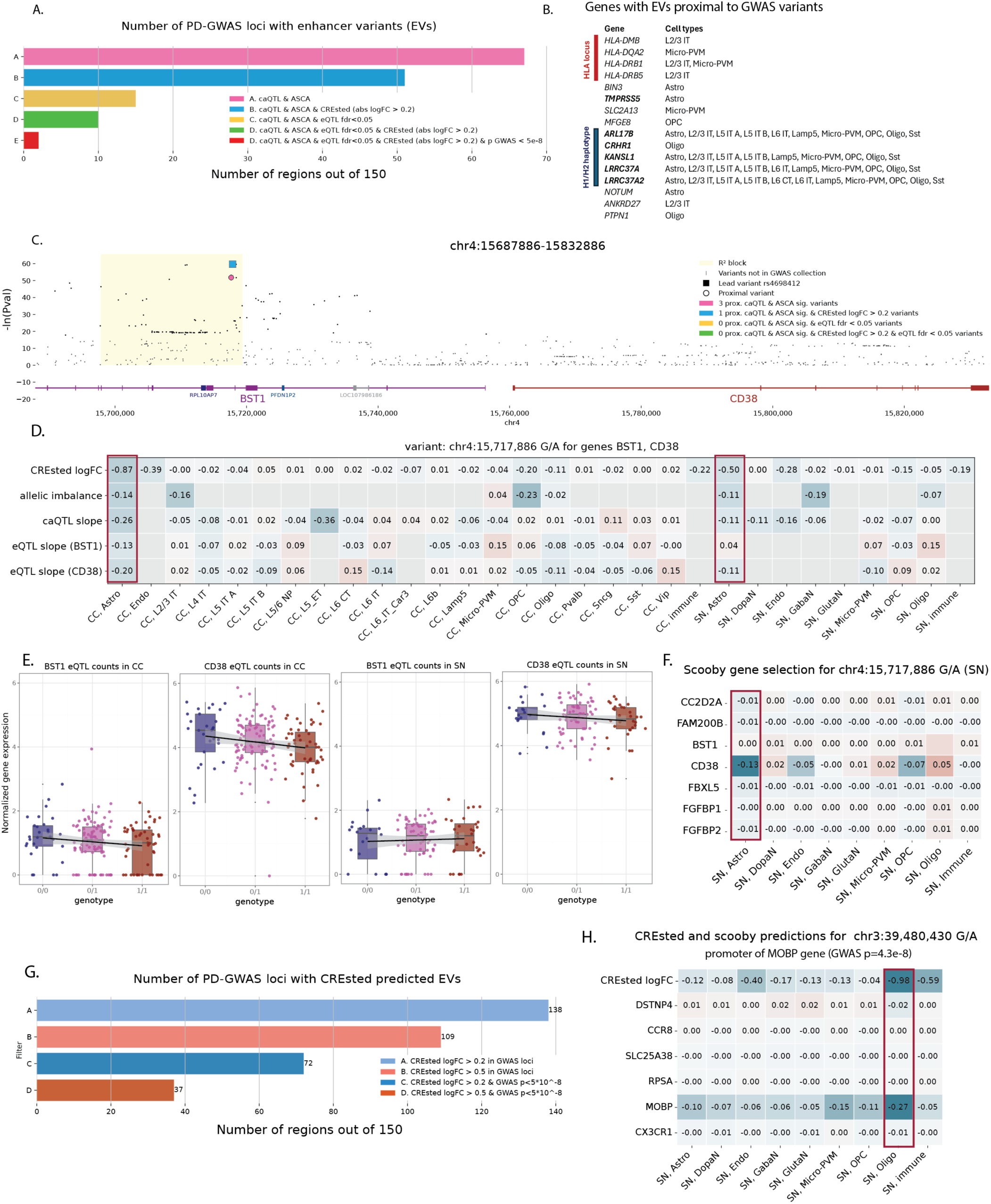
Regulatory variation at PD GWAS loci. **A.** Barplot of the number of PD GWAS loci (*n*=150) containing caQTL-ASCA variants; also predicted by CREsted; also an eQTL; or all of the above. **B.** Genes with enhancer variants (EV)s in PD GWAS loci from A (green bar), with the affected cell types. **C.** PD GWAS variants in locus chr4:15,687,886-15,832,886 near neurodegeneration-linked gene *CD38*. The R^2^-defined LD block (light yellow) contains 3 caQTL-ASCA variants, one of which – the block’s lead variant (rs4698412) – also has high CREsted logFC prediction. **D**. CREsted logFC, allelic imbalance, caQTL slope, and eQTL slopes for *BST1* and *CD38* across all tested cell types for variant chr4:15,717,886 G/A. Values in Astrocytes in CC and SN are highlighted by red boxes. **E.** Boxplots with per million normalized log1p transformed gene expression per donor of *CD38* and *BST1* in Astrocytes in both brain regions stratified by donor genotype. **F.** scooby logFC for all genes tested for variant chr3:39,480,430 G/A in all cell types in SN. **G.** Barplot of the number of PD GWAS loci containing CREsted predicted variants with logFC > 0.2 (blue) or logFC > 0.5 (red). Numbers of loci where CREsted predicted variants also have GWAS *p*-value <5e-8 are shown in dark. **H.** CREsted and scooby predicted logFC for chr3:39,480430 G/A variant in all tested SN cell types. scooby predictions are shown for all genes in the test window. The cell type with the highest CREsted and scooby effect predictions is highlighted (Oligodendrocytes, SN).

Although the majority of GWAS loci fall in noncoding regions, yet only a limited fraction colocalize with significant cis-eQTLs, highlighting the eQTL-GWAS paradox^51–53^. This gap likely reflects limited power, context specificity across tissues and cell states, and regulatory effects that are too small or transient to pass conventional eQTL significance thresholds^28,54^. Recent study on a big cohort of donors^29^ also demonstrated that GWAS genes had consistently smaller eQTL effect sizes compared to genes not associated with diseases suggesting more buffering mechanisms in essential genes. Based on all this, we next focused on EVs with strong model predictions but not passing eQTL significance thresholds. An interesting example is lead GWAS variant (rs4698412) located in *BST1* intron (Figure 7C), with the corresponding peak specifically accessible in astrocytes (Supplementary Figure 11A). Concordantly, this variant is predicted to disrupt chromatin accessibility in astrocytes by CREsted models confirmed by caQTL and ASCA in donors (Figure 7D). This variant has been assigned to *BST1* by proximity^10^, however *BST1* has bone marrow and intestine specific expression^55^. *CD38* is a paralogue of *BST1* located in the same locus with astrocyte specific expression and strong links to PD^56–59^. According to our analysis, both genes don’t pass eQTL significance thresholds, however *CD38* has high expression in donors’ astrocytes (Figure 7E) and its expression is predicted to be affected in SN astrocytes by scooby (Figure 7F) making it a strong putative target gene for rs4698412.

When focusing on CREsted predictions alone, we found 109 GWAS loci with strong putative EVs (CREsted logFC > 0.5), including 37 loci where predicted EVs passed global GWAS significance threshold (Figure 7G). Those include lead GWAS variant (rs356182) of *SNCA* locus with strong PD associations in multiple cohorts^60,10^, and an experimentally validated variant (chr12:40,161,320 C/T) shown to affect *LRRK2* expression in microglia^14^ (Supplementary Figure 11B-C). rs356182 was predicted to decrease chromatin accessibility specifically in SN dopaminergic neurons by disrupting FOXO3 binding motif (Supplementary Figure 11D), however we do not predict its effect on *SNCA* gene expression with scooby (Supplementary figure 11B), which agrees with recent experimental work confirming FOXO3 binding and suggesting an effect of this variant during development^15^. Another example is a GWAS SNP (p=4.3e-8) at chr3:39,480,430 (G/A), falling in the promoter of the oligodendrocyte-specific gene *MOBP*, and this variant is predicted to decrease its chromatin accessibility by disrupting flanking sequences in a SOX motif cluster (Supplementary Figure 11E). Although this variant does not have any observed effect in the donor snRNA-seq levels, it is predicted to affect *MOBP* gene expression by scooby (Figure 7H), making it an interesting candidate for further validation.

Overall, these analyses demonstrate that our framework can extract functional insights from PD GWAS loci by identifying putative effector variants and target genes even when direct eQTL support is weak or absent. By leveraging chromatin-based evidence and predictive models, we uncover cell-type-specific regulatory mechanisms that may underlie noncoding disease associations.

## Discussion

We created a unique resource to study functional cis-regulatory variation in the human brain, integrating long-read whole-genome sequencing (WGS) with matched single-cell multiomic atlases across a large cohort. By leveraging natural genetic variation as an *in vivo* mutagenesis system, we have moved beyond the limitations of *in vitro* assays, which often lack physiological context, to systematically map enhancer codes and variation thereof in the human brain and associated with Parkinson’s Disease (PD) risk.

High-confidence SNPs and indels from personal genomes were assessed for their correlation with chromatin accessibility through a combination of cohort-level variation (caQTL) and allelic imbalance (ASCA), yielding a set of cis-acting variants affecting cell type-specific regulatory elements. The high concordance between these orthogonal methods (Pearson’s *r* = 0.88) underscores the robustness in detecting how sequence variation correlates with chromatin accessibility. Furthermore, by incorporating bulk ONT methylation data, we demonstrated a strong relationship between increased chromatin accessibility and DNA hypomethylation, providing an additional layer of validation for these regulatory effects, and pinpointing alternative bulk measures that can serve as powerful proxies for single-cell caQTL/ASCA.

A central aspect of this work is the investigation of how sequence-to-function models identify and explain cis-regulatory variation, and how they can be used to overcome data limitations. Locally trained CREsted models to predict ATAC peak heights from the reference genome sequence can predict the functional impact of genetic variants directly from DNA sequence. Because these models are trained on the reference human genome and not donor-specific data, their high predictive accuracy (Spearman’s rho = 0.71–0.81 for caQTL-ASCA variants) suggests they have captured fundamental principles of the genomic cis-regulatory code. Once validated, we applied these models more broadly to score all variants, rather than only caQTL-ASCA, and demonstrate that they offer a significant advantage in overcoming the “statistical power gap” inherent in single-cell population studies. caQTL and ASCA mapping is limited by the number of cells in rare populations, such as dopaminergic neurons. S2F models on the other hand could predict variant effects in these disease-relevant cell types with high sensitivity, effectively prioritizing variants that lack sufficient power for standard statistical detection.

The “explainability gap” between GWAS loci and eQTLs remains a significant challenge in human genetics. Our results suggest that chromatin accessibility (caQTL-ASCA) and CREsted predictions may offer a more direct approach to interpret and prioritize GWAS signals than gene expression (eQTLs). This may be due to the buffering of essential genes or the fact that many regulatory variants exert subtle, condition-specific effects that are difficult to capture in postmortem tissue^29^. Nevertheless, about 10% of EV can be associated with eQTL-quantified variable gene expression of candidate target genes within a 1Mb sequence window. Across the two brain regions, this yielded 1,121 genes with variable gene expression linked to specific enhancers, including 233 genes with interpretable enhancer-level sequence changes (based on S2F models). We believe that this set of data-driven EV-gene pairs will form a ground truth benchmarking set to evaluate the capability of larger Borzoi/AlphaGenome-style foundation models to predict the effect of sequence variation on gene expression. We illustrated this idea using scooby-fine tuning of a base Borzoi model, which despite good performances on test data, remains limited in correctly predicting these EV-gene pairs (ca. 25% sensitivity), which aligns with recent reports^6,61,62^. Applying both EV-prediction and EV-gene pair prediction to GWAS loci linked to PD identifies and explains non-coding variation within 107 enhancers, of which 33 could be linked to putative target genes. Focusing on CREsted predicted EVs, we could further prioritize 103 enhancers affected by variants with strong GWAS signal, and some of those variants were further linked to target genes by scooby. This, for the first time, provides systematic gene-regulatory interpretations for multiple PD loci, affecting multiple cell types, including astrocytes, microglia and dopaminergic neurons. Key examples of genes with explained or predicted EVs include genes in the H1/H2 haplotype, *CD38*, and *SNCA*.

While our study revealed a considerable landscape of functional enhancer variants in the human brain, of which dozens in PD-linked GWAS loci, several avenues remain for future exploration to refine our understanding of genetic variation affecting regulatory logic across brain cell types and linked to diseases. Moving beyond the current reliance on pseudobulked samples toward emerging single-cell QTL methods might better resolve cell-state-specific effects^63^. Additionally, investigating the role of personal genomes and the combinatorial effects of multiple variants may significantly improve the accuracy of gene expression predictions, potentially uncovering regulatory interactions that single-variant analyses miss. Clearly, model improvements are needed to explain full gene loci at single-cell resolution^6^. Finally, the rich dataset of neurotypical controls and the CREsted and scooby models provided here serve as a valuable resource beyond the Parkinson’s field, offering a foundation to explore the functional basis of other neurological conditions, such as Alzheimer’s Disease, and broader phenotypic diversity in the human brain.

## Methods

### Human Postmortem Brain Tissue

The human sample collection and processing have been approved by the Ethics Committee Research UZ / KU Leuven (S64966). The postmortem human brain tissue has been obtained from three biobanks: the Edinburgh Brain Bank (University of Edinburgh, UK), the Banner Sun Health Research Institute^64^ (USA), and the Queen Square Brain Bank for Neurological Disorders (UCL Queen Square Institute of Neurology, UK). Frozen tissue from midbrain (SN), motor cortex, and anterior cingulate gyrus (CC) were used in this study and stored at −70 °C until processing. PD cases were selected to represent a mix of early and late disease stages, with a balanced ratio of males to females across both PD and control groups and similar age ranges between groups where possible. Donor and sample details are summarized in Supplementary Table 1.

### Tissue Sampling and Storage

For each region, tissue samples were isolated using disposable biopsy punches (Kai Medical, BPP-20F) on dry ice to maintain RNA and DNA integrity. Punches were collected in pre-chilled RNase/DNase-free tubes (Axygen, MCT-060-L-C) and stored at −70 °C until further processing. For single nuclei experiments, tissue punches of multiple donors were pooled together in a tube to scale up and reduce batch effects.

### Library preparation for long read sequencing

High Molecular Weight (HMW) DNA was extracted from cingulate gyrus tissue punches using either the NEB Monarch High Molecular Weight (HMW) DNA Extraction Kit for Tissue (New England Biolabs, T3060L) or the Nanobind Tissue Big DNA Kit (Circulomics, NB-900-701-01), according to the manufacturer’s instructions. DNA quantity was assessed using a Qubit fluorometer (Invitrogen). HMW DNA was sheared with 25 to 30 strokes of a 26-gauge blunt needle (SAI Infusion Technologies, 89134-164) prior to library preparation using the Ligation Sequencing Kit (Oxford Nanopore Technologies, SQK-LSK110 and SQK-LSK114), following the manufacturer’s instructions.

### Fiber-seq library preparation for long read sequencing

Fiber-seq requires specific processing prior to genomic HMW DNA extraction. A detailed step by step protocol is available from https://dx.doi.org/10.17504/protocols.io.j8nlkw54wl5r/v1. In brief, cingulate gyrus tissue punches were placed in a dounce homogenizer (KIMBLE® Dounce tissue grinder set, DWK885300-0001) on ice, and the tissue was gently homogenized in lysis buffer (10 mM Tris-HCl (pH 7.4), 10 mM NaCl, 0.1 % Bovine Serum Albumin, 0.5 mM Spermidine (pH 7.4) (Sigma-Aldrich, 85558), 0.1 % TWEEN® 20 (Sigma-Aldrich, P1379), 0.02 % Digitonin (Promega, G944A), 1× cOmplete Protease Inhibitor Cocktail (Roche, 04693132001), prepared in nuclease-free water) with 10 strokes of both the loose and tight pestles. After 5 minutes of incubation, we added 1000 µL wash buffer (10 mM Tris-HCl (pH 7.4), 10 mM NaCl, 0.1 % Bovine Serum Albumin, 0.5 mM Spermidine (pH 7.4), 0.1 % TWEEN® 20, prepared in nuclease-free water). The lysate was transferred to a 2 mL Protein LoBind tube and another 500 µL wash buffer was used to rinse the dounce and pestles after which it was added to the sample. The lysate was pelleted by centrifugation at 700 x g for 5 minutes at 4 °C after which the supernatant was discarded. Next, the nuclei were resuspended in 200 µL Hia5 labeling buffer (15 mM Tris-HCl (pH 8), 15 mM NaCl, 60 mM KCl, 1 mM EDTA (pH 8), 0.5 mM EGTA (pH 8), 0.1 % Bovine Serum Albumin, 0.5 mM Spermidine (pH 7.4), 0.8 mM S-adenosyl-methionine (NEB, B9003S), 5 µM Hia5 enzyme, prepared in nuclease-free water) and incubated in a ThermoMixer (Eppendorf) for 30 minutes at 37 °C and 900 rpm. The labelling was stopped by adding lysis buffer of the HMW DNA extraction kit. DNA extraction was performed as described above. Fiber-seq was also performed on two cell lines, NA12878 (RRID:CVCL_7526) and HBL (MM001) (RRID:CVCL_J075), to evaluate and finetune the models, respectively for SQK-LSK110 and SQK-LSK114. For these cell lines, the dounce homogenization was skipped, and cell pellets of one million cells were immediately lysed in the lysis buffer. All other steps remained the same.

### Long Read Sequencing

Sequencing of native DNA or 6mA-labeled DNA (Fiber-seq) was done on PromethION flow cells (Oxford Nanopore Technologies) for a minimum of 72 hours. Each flow cell was washed twice — each time followed by a reload of the library. Samples with low coverage or low read quality were re-sequenced on a second flow cell. Sequencing was performed using SQK-LSK110 (*n* = 51) or SQK-LSK114 (*n* = 127) from high molecular weight DNA extracted from CC (*n* = 168), gut (*n* = 16), or SN (*n* = 4). HMW DNA extraction failed for 2 donors due to sample quality issues of both brain regions.

### WGS data processing

#### Fiber-seq model training

In short, Fiber-seq introduces exogenous m6A methylation on accessible adenosines, providing information about open chromatin in long-read WGS. Dense exogenous labelling of adenosines with Hia5 enzyme during Fiber-seq can increase basecalling error rates in Nanopore sequencing. To improve basecalling accuracy and support high-quality variant calling, we finetuned basecalling models provided by ONT. Specifically, Bonito (0.5.0; RRID:SCR_028115) was used to finetune Guppy (RRID:SCR_023196) basecalling model dna_r9.4.1@v3.3. Reads, both passing and failing quality checks from four separate ONT Fiber-Seq runs were subsampled and used as training data. The model was evaluated by basecalling ONT Fiber-Seq data from Genome In A Bottle (GIAB) sample NA12878. Data was aligned to hg38 without alt contigs using minimap2^65^ (2.24; RRID:SCR_018550) and small variants were called using Clair3^66^ (0.1-r12; RRID:SCR_026063) using the /opt/models/r941_prom_sup_g5014 model. Precision, recall and F1 scores for SNP and indel calling were calculated based on GIAB HG001/NA12878 v4.2.1 benchmark calls in select regions using hap.py (0.3.12, docker image pkrusche/hap.py:latest; variant calls: https://ftp-trace.ncbi.nlm.nih.gov/ReferenceSamples/giab/release/NA12878_HG001/NISTv4.2.1/GRCh38/).

For SQK-LSK114 4kHz Fiber-seq data: The pre-trained dna_r10.4.1_e8.2_400bps_sup@v4.1.0 model was fine-tuned using Bonito (0.7.2) on ONT Fiber-Seq data generated from the HBL (MM001) cell line. The model was fine-tuned using 250,000 randomly selected reads for one epoch and using learning rate of 5e-4. Evaluation was done on a held-out reads test set, achieving mean and median accuracies of 98.844% and 98.979%, respectively.

#### NanoWGS pipeline

Raw long-read nanopore WGS data was processed using an in-house developed Nextflow pipeline titled NanoWGS (https://github.com/AlexanRNA/nanowgs/releases/tag/v0.0.2 / zenodo: 10.5281/zenodo.13385064, RRID:SCR_028113). Pipeline steps are detailed below. Due to meaningful updates to ONT chemistries and software during this project, subsets of samples have been processed with different tool versions, in particular Dorado and Guppy (but also SAMtools and minimap2). The specific versions used for each sample are collated in Supplementary table 2.

#### Basecalling

After sequencing, fast5 (older) or pod5 (more recent) files were basecalled using Guppy or Dorado (see Supplementary table 2). To improve basecalling speeds with Dorado, fast5 files were typically first converted to pod5 using pod5tools (0.2.0; RRID:SCR_028166).

#### Alignment

In case of Dorado basecalling, unaligned bams were first converted to fastq using SAMtools^67^ (RRID:SCR_002105) command ‘samtools fastq-T ‘*’’. This step was not necessary for Guppy, as fastq files were output by default. Reads were aligned to the T2T-CHM13v2.0 reference genome^68^ using minimap2^65^. The coverage and other basic quality statistics were computed using Cramino^69^ (0.14.5; RRID:SCR_028160) and mosdepth^70^ (0.3.3; RRID:SCR_018929).

#### Variant calling

SNPs and indels were called using Clair3^66^ (1.0.9) using the relevant ONT model, ‘--platform=“ont”’, and default parameters. Variants were filtered using BCFtools^67^ (v1.20; RRID:SCR_005227) to keep PASS variants < 50 bps and alternative alleles were trimmed with--trim-alt-alleles. SV calling was performed using Sniffles2^71^ (2.2; RRID:SCR_017619) specifying a minimum variant size of 50 bps and providing tandem repeat annotations from http://t2t.gi.ucsc.edu/chm13/dev/old.t2t-chm13-v2.0/trf/trf.bigBed in--tandem-repeats parameter. BCFtools (1.20) was used to retain PASS SV calls only.

#### Phasing

Phasing of SNPs, indels and SVs was done using the LongPhase^72^ ‘longphase phase’ command (1.7.3; RRID:SCR_028162) after which reads in the BAMs were tagged using ‘longphase tag’. Finally, BAM files were compressed into CRAMs using SAMtools (1.20).

#### CpG Methylation

CpG methylation info was encoded in BAMs for samples basecalled with Dorado. We did not call methylation states for samples basecalled with Guppy. Based on the model used (Supplementary table 2), a subset of samples have CH methylation available as well. To avoid confounding by tissue-of-origin effects, we only included samples from CC in methylation analyses (*n*=125). Methylation rates at CpG sites were extracted from the processed CRAM files using Modkit (v0.4.3; RRID:SCR_028163) with the following command ‘modkit pileup $input_cram $output_bed --ref $fasta --cpg --combine-strands’. To visualize methylation data, we used methylartist^73^ (v1.5.3; RRID:SCR_028164)

#### Population small variant calling

Individual phased VCF files produced by Clair3 and phased by LongPhase were first combined with BCFtools^67^ v1.21 command “bcftools merge --missing-to-ref --merge all”. This combines multiallelic records if needed by merging SNP and INDEL records, while setting all unseen alleles to the reference one. To produce input genotype files for single-cell donor demultiplexing (see below), only biallelic SNPs were retained with “bcftools +remove-overlaps” and “bcftools view --types snps”.

#### Population structural variant calling

Population structural variant calling was done using Sniffles2 (2.6.0) with default parameters apart from ‘--combine-max-inmemory-results 200’ and ‘--combine-pctseq 0.5’. Five samples were excluded from population SV calling based on either outlier high numbers of reported inversions (ASA_003B, ASA_106B), outlier low numbers of deletions (ASA_003B, ASA_004B, ASA_040B, ASA_106B), or based on manual inspection (ASA_025B). The population VCF was then filtered using BCFtools (v1.21) to remove SVs with AF=0.0 and AF=1.0.

#### Whole genome sample identity quality check

To ensure there were no sample swaps or inconsistencies, we cross-checked WGS-derived sample sex with biobank metadata. We also used somalier^74^ (v0.2.19; RRID:SCR_028167) to estimate sample relatedness and ancestry. Donor deconvolution in single cell data (described in ‘Donor demultiplexing’ below) served as an additional check, since we could check if we observed the expected donors in each donor pool. No inconsistencies were observed and no samples were excluded based on these checks.

#### Clinvar overlap

To identify donors who potentially have familial PD or carry known highly penetrant mutations but are misdiagnosed as idiopathic, we overlapped each patient’s SNP calls with Clinvar using bcftools isec (1.19). SNPs were retained if the VCF “INFO” field contained “Parkinson” and “pathogenic” and did not contain “benign” (reported in Supplementary table 9).

#### Library Preparation for Single Cell Short Read Sequencing

For all single nuclei experiments, the biopsies were taken from-70 °C storage and kept on dry ice until the tissue was equilibrated in the homogenization buffer. All buffers were made fresh and stored on ice. Frozen tissue biopsies were homogenized and lysed using a pre-chilled 1 ml Dounce homogenizer (Kimble) containing 750 µL cold homogenization buffer and by stroking up and down 10 times with both the loose pestle and the tight pestle.

#### Nuclei preparation for 10x Genomics Multiomic assay

Nuclei for the 10x Genomics Multiomic assay were isolated as described by the following protocol: https://dx.doi.org/10.17504/protocols.io.n2bvj3yqnlk5/v1. In brief, the tissue was homogenized and lysed in Homogenization Buffer (10 mM Tris-HCl (pH 7.5), 146 mM NaCl, 1 mM CaCl_2_, 21 mM MgCl_2_, 1% BSA (Gibco, 15260037), 25 mM KCl, 250 mM sucrose, 0.03% TWEEN® 20, 0.03% IGEPAL® CA-630 (Sigma-Aldrich, I8896), 1 mM dithiothreitol, 1× cOmplete protease inhibitor (Roche), and 1 U/µL RNasin® Plus Ribonuclease Inhibitor (Promega, N261B)). After lysis the nuclei were first filtered through a 70 µm EASYstrainer (Greiner, 542070), mixed with 1 mL of Wash Buffer 1 (WB1) (10 mM Tris-HCl (pH 7.5), 146 mM NaCl, 1 mM CaCl_2_, 21 mM MgCl_2_, 1% BSA, 25 mM KCl, 250 mM sucrose, 1 mM dithiothreitol, 1× cOmplete protease inhibitor (Roche), and 1 U/µL RNasin® Plus Ribonuclease Inhibitor (Promega)) and centrifuged (500 rcf) in a swinging bucket for 5 minutes at 4°C. The supernatant was removed and the nuclei were resuspended in a total volume of 520 µL of WB1 and then gently mixed with 520 µL of Gradient Medium (10 mM Tris-HCl (pH 7.5), 1 mM CaCl_2_, 5 mM MgCl_2_, 50% iodixanol (Sigma-Aldrich, D1556), 75 mM sucrose, 1 mM dithiothreitol, 0.5× cOmplete protease inhibitor (Roche), and 1 U/µL RNasin® Plus Ribonuclease Inhibitor (Promega)). Then the sample was layered upon 770 µL of iodixanol cushion (31 mM Tris-HCl (pH 8.0), 15.5 mM MgCl_2_, 77.5 mM KCl, 129.2 mM sucrose, 29% iodixanol and 0.3 U/µL RNasin® Plus Ribonuclease Inhibitor (Promega)). Density gradient centrifugation for 20 minutes in a swinging bucket (3,000 rcf) was used to remove debris. Afterwards, nuclei were further permeabilized with a lysis buffer (10 mM Tris-HCl (pH 7.5), 10 mM NaCl, 3 mM MgCl_2_, 0.5% BSA, 0.01% TWEEN® 20, 0.01% IGEPAL® CA-630, 0.001 % Digitonin, 0.5 mM dithiothreitol, and 0.5 U/µL RNasin® Plus Ribonuclease Inhibitor (Promega)), washed with Wash Buffer 2 (10 mM Tris-HCl (pH 7.5), 10 mM NaCl, 3 mM MgCl_2_, 1% BSA, 0.01% TWEEN® 20, 1 mM dithiothreitol, and 1 U/µL RNasin® Plus Ribonuclease Inhibitor (Promega)), and resuspended in Diluted Nuclei Buffer (1× Nuclei Buffer (10x Genomics, PN-2000207), 1 mM dithiothreitol, Protector Rnase Inhibitor (Sigma-Aldrich, 3335402001)) as described by the 10x Genomics’ User Guide (10x Genomics, CG000338).

#### Nuclei preparation for 10x Genomics snRNA-seq and Parse Biosciences

Nuclei that were used for 10x Genomics Chromium Single Cell 3ʹ GEM v3 or Parse Biosciences Evercode™ v1 WT were purified with the following protocol: https://dx.doi.org/10.17504/protocols.io.5qpvo1rddg4o/v1. In brief, the tissue was homogenized and lysed in Homogenization Buffer with Tween 20 (10 mM Tris-HCl (pH 7.5), 146 mM NaCl, 1 mM CaCl_2_, 21 mM MgCl_2_, 0.01% BSA, 25 mM KCl, 250 mM sucrose, 0.03% TWEEN® 20, 1 mM dithiothreitol, 0.5× cOmplete protease inhibitor (Roche), and 0.5 U/µL RNasin® Plus Ribonuclease Inhibitor (Promega)). After lysis the nuclei were filtered through a 70 µm EASYstrainer (Greiner), and centrifuged (500 rcf) in a swinging bucket for 5 minutes at 4°C. The supernatant was removed and the nuclei were resuspended in a total volume of 520 µL of Homogenization Buffer without Tween 20 (10 mM Tris-HCl (pH 7.5), 146 mM NaCl, 1 mM CaCl_2_, 21 mM MgCl_2_, 0.01% BSA, 25 mM KCl, 250 mM sucrose, 1 mM dithiothreitol, 0.2× cOmplete protease inhibitor (Roche), and 0.2 U/µL RNasin® Plus Ribonuclease Inhibitor (Promega)) and then gently mixed with 520 µL of Gradient Medium (10 mM Tris-HCl (pH 7.5), 1 mM CaCl_2_, 5 mM MgCl_2_, 50% iodixanol, 75 mM sucrose, 1 mM dithiothreitol, 0.5× cOmplete protease inhibitor, and 0.5 U/µL RNasin® Plus Ribonuclease Inhibitor (Promega)). Then the sample was layered upon 770 µL of iodixanol cushion (31 mM Tris-HCl (pH 8.0), 15.5 mM MgCl_2_, 77.5 mM KCl, 129.2 mM sucrose, and 29% iodixanol). Density gradient centrifugation for 20 minutes in a swinging bucket (3,000 rcf) was used to remove debris. Afterwards, the supernatant was removed, and the nuclei were resuspended and washed with Resuspension Buffer (1× PBS, 0.75 % BSA, 1 U/µL RNasin® Plus Ribonuclease Inhibitor (Promega)) by centrifugation at 350 rcf for 5 minutes. Supernatant was again removed, and the nuclei were resuspended in Resuspension Buffer. These nuclei were either processed with Chromium Single Cell 3ʹ GEM v3 as described by the 10x Genomics’ User Guide (10x Genomics, CG000183) or Parse Biosciences Evercode™ v1 WT (dx.doi.org/10.17504/protocols.io.8epv55xwdv1b/v1).

#### Nuclei preparation for 10x Genomics snATAC-seq and scaled snATAC-seq

We used the following protocol to purify nuclei for snATAC-seq and scaled snATAC-seq: https://dx.doi.org/10.17504/protocols.io.14egnrb7ql5d/v1. In brief, the tissue was homogenized and lysed in Homogenization Buffer plus detergents (10 mM Tris-HCl (pH 7.5), 10 mM NaCl, 3 mM MgCl_2_, 0.04 % BSA, 250 mM EDTA, 320 mM sucrose, 0.01 % TWEEN® 20, 0.05% IGEPAL® CA-630, 0.0025 % Digitonin,1 mM dithiothreitol, and 1× cOmplete protease inhibitor (Roche)). After lysis the nuclei were filtered through a 70 µm EASYstrainer (Greiner), and centrifuged (500 rcf) in a swinging bucket for 5 minutes at 4°C. The supernatant was removed and the nuclei were resuspended in a total volume of 520 µL of Homogenization Buffer without detergents (10 mM Tris-HCl (pH 7.5), 10 mM NaCl, 3 mM MgCl_2_, 0.04 % BSA, 250 mM EDTA, 320 mM sucrose, 1 mM dithiothreitol, and 1× cOmplete protease inhibitor (Roche)) and then gently mixed with 520 µL of Gradient Medium (10 mM Tris-HCl (pH 7.5), 5 mM CaCl_2_, 3 mM MgCl_2_, 50% iodixanol, 0.04 % BSA, 1 mM dithiothreitol, and 0.2× cOmplete protease inhibitor). Then the sample was layered upon 770 µL of iodixanol cushion (31 mM Tris-HCl (pH 8.0), 15.5 mM MgCl_2_, 77.5 mM KCl, 129.2 mM sucrose, and 29% iodixanol). Density gradient centrifugation for 20 minutes in a swinging bucket (3,000 rcf) was used to remove debris. Afterwards, the supernatant was removed, and the nuclei were resuspended and washed with Resuspension Buffer (1× Nuclei Buffer (10x Genomics, PN-2000207), 1 % BSA) by centrifugation at 350 rcf for 5 minutes. Supernatant was again removed, and the nuclei were resuspended in Resuspension Buffer. Samples were either immediately processed with Chromium Next GEM Single Cell ATAC v2 (10x Genomics, PN-1000390), HyDrop-ATAC v2 (dx.doi.org/10.17504/protocols.io.x54v97mmpg3e/v1), or first pre-indexed with barcoded Tn5, which is described in more detail in the following protocols: Scale Biosciences scATAC Pre-Indexing (https://www.protocols.io/view/scatac-pre-indexing-kit-10x-scatac-seq-n92ld5bj8v5b/v1), or an in-house developed protocol for scaled snATAC-seq ( dx.doi.org/10.17504/protocols.io.4r3l2z24ql1y/v1, dx.doi.org/10.17504/protocols.io.j8nlk1w1wg5r/v1).

#### Short Read Sequencing

Sequencing libraries from snRNA-seq and snATAC-seq were sequenced on the following Illumina platforms: NextSeq500, NextSeq2000, NovaSeq6000 v1.5, and Illumina NovaSeqX. We always used at least the minimum of read cycles that were required for the specific assay, as per manufacturer’s directions.

### Single nuclei RNA-seq data processing

#### 10X RNA and 10X multiome RNA-seq data processing

First, the sequencing data has been processed by CellRanger (8.0.1; RRID:SCR_017344) using custom CHM13v2 index using the following command: ‘cellranger count --libraries=“${libraries}”--transcriptome=“${cellranger_transcriptome_dir}” --create-bam=“true” --expect-cells=“1 --include-introns=true --chemistry=ARC-v1’. Next, we performed quality check for each multiome lane separately using Scanpy (1.10.4; RRID:SCR_018139). First, nuclei with less than 200 genes expressed and genes expressed in fewer than 3 cells were filtered out. We used median absolute deviation (MAD) as described in Heumos et al.^75^ to identify outlier nuclei. We excluded all nuclei with log counts, log1p_n_genes_by_counts or pct_counts_in_top_20_genes with five times over MAD. Nuclei with mitochondrial counts above 12%, cells with over 75,000 counts and nuclei with over 10,000 genes detected were also filtered out. The doublet rate was estimated using Scrublet (RRID:SCR_018098). The data from all the multiomes was combined into a single object, normalized, top 3,000 highly variable genes were selected and PCA, neighborhood graph, Leiden clustering, UMAP and t-SNE were performed. After donor demultiplexing (described in ‘Donor demultiplexing’), the information on donor assignments was added and used to add region information as well. Afterwards, nuclei objects were separated into two anndata objects, one for SN and one for CC together with motor cortex. These datasets were then reclustered, and new UMAP projection coordinates were calculated. Using Leiden clusters at resolution 4.5, we identified all clusters with doublet assignments for ≥ 30% of nuclei. Any nuclei that fell into these clusters, had vireo assignment (described in Donor demultiplexing section) of ‘doublet’ or ‘unassigned’, or were predicted to be a doublet by Scrublet were removed from further analysis. In both datasets, we additionally identified clusters with low *MALAT1* expression, which can be used as a proxy for low nuclear fraction droplets^76^ and removed them. Afterwards, data was reclustered.

#### ParseBiosciences snRNA-seq data processing

For the ParseBio RNA, we used ParseBiosciences’ Split-pipe pipeline (1.3.1; Proprietary Software) with CHM13v2 index to obtain cell x gene count matrix. We performed processing for the two ParseBiosciences experiments separately. For 100k cells kit, we first processed all sequencing sublibraries with the following command: ‘split-pipe --mode all --chemistry v1 --kit WT’, whereas for 1M kit, we used the same command but with parameter ‘--kit WT_mega’. For both 1M and 100k kit, the sublibraries were combined into one object using ‘split-pipe --mode comb’. Afterwards, data was loaded in Scanpy (1.10.4). For 100k kit (PB-001 in this manuscript), we filtered out nuclei with less than 100 unique UMIs, less than 200 genes and genes expressed in fewer than 3 cells. For 1M kit (PB-002), we filtered out nuclei with fewer than 200 unique UMIs and genes expressed in fewer than 5 cells. For both PB-001 and PB-002, 5*MAD of log1p_total_counts, log1p_n_genes_by_counts, pct_counts_in_top_20_genes, pct_counts_mt and nuclei with over 12.5% mitochondrial counts were excluded from downstream analysis. For PB-002, which contained data from both CC and SN, the data was then divided based on the region it was originating. For ParseBiosciences data, Scrublet was not used to estimate doublets. Instead, the clusters at Leiden resolution of 2.5 for PB-001 and of 4.5 for PB-002 (for both CC and SN data) that had doublet fraction of ≥30%, or nuclei that had vireo assignments of ‘doublet’ or ‘unassigned’ were removed from the downstream analysis. The data was then reclustered again.

#### Single nuclei RNA cell type annotation

To annotate the cell types, we trained scANVI models^21^ in Scanpy (1.9.1) with scvi-tools (0.17.4; RRID:SCR_026673). For CC, we trained scANVI model using published human motor cortex data^77^.This model was trained with top 2,000 highly variable genes and using external_donor_name_label as batch key. The model was trained with the following parameters: use_layer_norm=“both”, use_batch_norm=“none”, encode_covariates=True, dropout_rate=0.2 and n_layers=2. For SN, scANVI model trained on published SN dataset^78^ with donor ID as batch key and top 2,000 variable genes was used. Default parameters were used. We ran the models separately on 10X SN data, 10X cingulate gyrus data, PB-001 and PB-002 data. After the automatic cell type annotation, the annotations were manually checked and adjusted based on the known curated cell type markers. Additionally, since motor cortex lacks L4 IT neurons, these were manually added to the cell type annotation based on markers genes (*RORB*+, *CUX2*+, *GRIN3A*-).

During analysis of CC data, based on clustering and marker gene expression, a donor containing striatum brain region contamination instead of CC was identified and excluded from the downstream analysis (donor ID: ASA_105).

#### Single nuclei RNA integration and annotation adjustments

To integrate single nuclei RNA data from 10X and ParseBiosciences, we used combination of scVI model^79^ using either 10X, 100k (PB-001) or 1M ParseBiosciences (PB-002) kit as batch key followed by scANVI^21^ model using cell type annotation as label key (scvi-tools 1.2.2.post2). After integration, we again confirmed the cell type annotations with marker genes for both regions. For CC data, using majority voting on Leiden clustering of resolution 2.5, we assured there are no discrepant cell type annotations within the same Leiden cluster. For both CC and SN, immune cells were added to annotation based on marker gene expression (*RUNX1*, *IL7R*, *CD8A*), the neuron annotations were also adjusted using cell type markers for dopaminergic (*TH*, *SLC6A3*, *SLC18A2*), gabaergic (*GAD1*, *GAD2*) and glutamatergic (*SLC17A6*, *SLC17A7*) neurons.

#### Differential gene expression

Differential gene expression was performed on per cell type and donor pseudobulked data using dreamlet^23^ (1.6.0; RRID:SCR_028168). Pseudobulks for each cell type were created for donors with minimum 5 cells and 5 counts. The healthy controls were contrasted against patients affected by PD. For each tested gene and cell cluster, the following mixed model was fitted to perform differential gene expression: expression ∼ age_at_collection + sex + biobank_name + batch + postmortem interval (PMI) + PD. FDR correction was performed separately for each brain region.

### Single nuclei ATAC-seq data processing

#### Preprocessing

Cell barcodes were extracted from FASTQ files and corrected with single_cell_toolkit (RRID:SCR_028169; commit 6e31c3b: 10x snATAC and 10x snMultiome ATAC with correct_barcode_from_fastq.sh, HyDrop snATAC v2 with extract_and_correct_hydrop_atac_barcode_from_fastq.sh, scaled snATAC with 10x snATAC barcodes or HyDrop snATAC barcodes with extract_and_correct_scalebio_atac_barcode_from_fastq.sh) and added to read names as SAM tags and values (Cellular barcode: CR:Z:sequence+, Phred quality of the cellular barcode sequence in the CR tag: CY:Z:qualities+, Cell identifier, consisting of the corrected cellular barcode sequence and library name CB:Z:str) of trimmed reads (Trim Galore v0.6.7; RRID:SCR_011847). Trimmed reads (of the same library) with corrected cellular barcodes were mapped with bwa-mem2 (v2.2.1; RRID:SCR_022192) with-C option to add SAM tags from read name to output BAM file to reference T2T-CHM13v2.0 sorted by genomic coordinate and merged using SAMtools (v1.19.1; RRID:SCR_016366) sort and merge. After merging BAM files of the same library, fragment files were made with create_fragments_file of single_cell_toolkit.

#### Quality control

For quality control and downstream processing of fragment files, pycisTopic^22^ (v2.0a0; RRID:SCR_026618) was used, and separate analyses were conducted for CC and SN. For each brain region, two subsequent analyses were performed; first, an exploratory analysis on a subset of the snATAC samples with preliminary annotations, for enriching the sets of consensus regions, followed by a final, in-depth analysis, integrating the complete snATAC dataset. In both cases, barcodes with TSS enrichment below 5 and less than 1,000 unique fragments in peaks were excluded. For the preliminary analysis, consensus regions from prior analyses were used, where pycisTopic was run on subsets of single nuclei multiome samples using the hg38 (GRCh38) reference genome. Consensus regions with 500 nucleotide length were generated and converted to T2T-CHM13v2.0 using UCSC liftOver^80^, which was also used for obtaining T2T-CHM13v2.0 TSS and blacklisted regions. For the final analysis, consensus regions obtained from the preliminary analysis were used.

#### Topic modeling and cell type analysis

Topic modeling was performed and respectively 100 and 60 topics were selected for CC and SN in the preliminary analysis. In the final analysis, 110 topics were selected for CC and 80 topics for SN. For obtaining the preliminary annotation, a majority voting strategy depending on RNA annotations of multiome cells was applied. Clusters from Leiden clustering with resolution 3.5 were assigned to a cell type if over 50% for CC and 70% for SN were of that cell type. Clusters not meeting these criteria and clusters with doublet label were excluded from the analysis. For the final analysis, a more stringent threshold of 70% and with resolution 9.5 for CC and resolution 5.5 for SN was used. For CC, a cluster of SNCG interneurons is present using a 60% threshold but is lost using the 70% threshold. After careful consideration and discussion, this cluster was annotated manually, given the strong indication of SNCG interneuron presence. Additionally, cells from one donor were excluded from the CC analysis, as was done in RNA (donor ID: ASA_105). For visualization, the first two dimensions of UMAP dimensionality reduction were plotted. Consensus peaks from the respective final annotations were obtained using the MACS2 (v2.2.9.1; RRID:SCR_013291) algorithm with default parameters and excluding blacklisted regions, as implemented in pycisTopic. From the final cell type annotations, differentially accessible regions (DARs) per cell type were calculated with default pycisTopic settings, and consensus regions were obtained as described above, resulting in 1,701,443 and 1,216,009 consensus regions for CC and SN, respectively. Pseudobulked snATAC coverage tracks were generated in BigWig file format and these tracks were used further for training peak regression CREsted models.

#### Donor demultiplexing

We used cellsnp-lite^81^ (v1.2.3; RRID:SCR_025515) and vireo^82^ (v0.5.9; RRID:SCR_027463) to genotype cells and assign them to their respective donors. For every single cell experiment pool, we subset the population BCF file to contain only SNPs of the donors present in the given pool. Additionally, for RNA libraries, we only kept SNPs that are covering gene body regions and for ATAC, we kept SNPs within consensus peaks. For 10X RNA and 10X multiome RNA, we ran the demultiplexing for cell-associated barcodes as called by cellranger (ie cell barcodes present in filtered_feature_bc_matrix). For ParseBiosciences RNA, we equally ran demultiplexing only on cell-associated cell barcodes. For ATAC libraries, only select barcodes passing QC, namely >=1000 fragments and TSS score of >=5.0, were assigned genotypes. Additionally, for ATAC libraries, the following parameter was added in cellsnp-lite ‘--UMItag None’.

### eQTL analysis

#### Plink

For eQTL analysis: Plink2^83^ (v2.0.0-a.6.1LM; RRID:SCR_001757) has been used to preprocess data for QTL analysis. For both SNPs and indels, variants with MAF < 0.01, --hwe < 1e-5 (Hardy-Weinberg equilibrium test) and more than two alleles were filtered out. For meQTL analysis, the same filters were applied as for eQTL, but since the analysis was performed on a subset of 125 donors, allele frequency was recalculated for the variants.

#### eQTL data preprocessing

eQTL analysis has been performed on pseudobulked samples. Donor-specific pseudobulks were prepared from single cell RNA data using pertpy python package^84^ (0.9.5; RRID:SCR_028170). To create a pseudobulk, a donor had to have ≥50 cells and ≥1,000 counts across those cells. Additionally, only cell types with least 30 donors were kept for analysis. Raw counts were then counts per million normalized and log1p transformed. Only protein coding genes were kept in the final donor x gene expression matrix.

#### eQTL mapping

eQTL was performed using tensorQTL^26^ package (1.0.10; RRID:SCR_028171) with the cis.map_nominal function. The QTL analysis was run separately for each cell type and for SNPs and for indels. We tested all protein coding genes and we used age, biobank, sex, sequencing method (10X, ParseBio or mix of both) and first five genotype principal components from Plink2 as covariates. Variants within ±1Mb window from the start and end of gene body were tested. FDR correction of 5% was performed on the set of all variants across all cell types.

### Chromatin accessibility QTL (caQTL) and allele specific chromatin accessibility (ASCA) analysis

#### caQTL data preprocessing

For the caQTL analysis, new fragment count matrices were generated using the cell type annotations and consensus regions obtained from the final datasets. Cells were grouped per cell type and donor, and groups with less than 50 cells were excluded. As in the eQTL data processing, cell types with less than 30 donors were excluded. Next, fragment counts were summed over all cells per combination, followed by counts per million normalization and log-transformation (using Numpy’s log1p).

#### caQTL mapping

As in the eQTL analysis, caQTL was performed using the tensorQTL package (1.0.10) with the cis.map_nominal function and was run separately for each cell type. Variants tested were required to fall within the region for SNPs and within ± 200 base pairs from the start or end of the region for Indels. Covariates used are age, biobank, sex, broad sequencing method (multiome, snATAC or scaled snATAC), first five genotype principal components from Plink2 and the first 30 principal components from a PCA analysis per cell type, on the aggregated per donor fragment counts.

#### Mappability filter for the ASCA analysis

For the ASCA analysis, we reprocessed snATAC-data to obtain allele-specific read counts of CC and SN consensus peaks per donor and cell type. First, BAM files were split per donor using demultiplexed cell barcodes, as described above. To account for potential mappability issues that can arise when reads containing different alleles are mapped to the reference genome, we applied WASP mappability filter^27^ (RRID:SCR_025497) using indel-aware implementation of the WASP pipeline^85^, which was further adapted to account for cell barcode information and phasing blocks (https:/github.com/OlgaSigalova/wasp_indel_v1.0). For each donor, mapped reads in the BAM file were overlapped with the personal VCF file, and all reads overlapping SNPs and Indels were filtered out and converted back in FASTQ file. Alleles in those reads were substituted with all possible allele combinations from the donor VCF file (up to 64 different allelic combinations per read) and mapped back to the T2T-CHM13v2.0 reference genome with the same options as specified above. If the read with all possible allelic combinations mapped to the same genomic location with a mapping quality above 10 it was retained, otherwise discarded as causing potential mappability issues. Resulting filtered BAM files were then split per cell type for subsequent quantification of allelic counts.

#### Getting allele specific counts per ATAC-seq peak

We quantified allelic counts for all variants falling within 500 bp windows of consensus peak sets of CC and SN (referred to as *test variants*). For each donor and cell type, we obtained number of reads containing REF and ALT allele for each test variant. These counts were then aggregated per peak for each test variant, by summing counts from all heterozygous variants in the peak and accounting for the phasing information with the test variant.

#### ASCA mapping

The allele-specific part of combined haplotype test (AS-CHT) from WASP suite was used to define variants associated with allelic imbalance across donors. The test was run separately for each cell type of CC and SN, including all test variants on autosomes with at least 5 heterozygous donors and at least 50 allele-specific reads in total (across donors). Allelic imbalance metric was defined as 𝛽/(𝛼 + 𝛽), where 𝛽 and 𝛼 are expected read counts estimated by AS-CHT.

#### Final set of variants and multiple testing correction

In the final set, we included variants quantified by both caQTL and ASCA (requiring at least 30 donors with at least 50 cells for a given cell type, including at least 5 heterozygous donors with minimum 50 allele-specific reads for a given ATAC peak), resulting in 2.2M variants (2M SNPs and 200K indels) in 1.5M consensus peaks. P-values from both caQTL and ASCA analyses were corrected using Benjamini-Hochberg correction^86^ (FDR) on the full set of tests combined (all variants tested in all cell types of CC and SN, 10.7M in total)

#### meQTL analysis

For methylation QTL analysis, we used the ATAC consensus peak set from CC as the phenotypes to be tested. In short, we took Modkit generated bed files of 125 donors for whom we had methylation data available and overlapped them with ATAC bins. We kept the bins that had information on methylation for at least 5 Cs in the 500 bp window. The methylation signal was then averaged across the 500 bp window. Only bins with methylation information across all donors were tested. Variants within 10,000 bp window of the methylation bin were tested. Donor sex, age, biobank name, basecalling model and first 5 genotype principal components from Plink2 were used as covariates. QTL analysis was performed using tensorQTL package (1.0.10) with the cis.map_nominal function for SNPs and indels separately.

#### Crested models

We used CREsted software package^4^ (RRID:SCR_026617) to model cell type specific enhancers and predict effects of genetic variation on chromatin accessibility from the genome sequence. We trained convolutional neural network models (Dilated CNN architecture, regression models, supplementary table) that take as input 2114bp DNA sequences (centered on consensus peak summits) and as an output predict a scalar of mean peak accessibilities across cell types. Separate models were trained for CC (21 cell types) and SN (9 cell types) on the corresponding consensus peak sets for these two brain regions. Peak heights were calculated by taking mean coverage in the central 1000bp sequences of the resized peaks and normalized across cell types with min-max normalization using top 3% of constitutive peaks per cell type (default). Train-validation-test set split was done by chromosome, with chr2 and chr22 in the validation and chr4 and chr9 in the test set, while the rest of the data was using for training (1.36M peaks in CC, 977K peaks in SN). Trained models (35 and 21 epochs for CC and SN, respectively) were then finetuned on the cell-type specific regions (regions with a Gini index 1 standard deviation above the mean across all regions, 275K and 194K peaks for CC and SN, respectively), with the same train-test-validation split as above.

CREsted models were used to score SNP and indel effects on chromatin accessibility using a peak-based in silico mutagenesis (ISM) approach. For each variant, the associated peak was resized to the model input length, and the alternative allele was introduced into the reference sequence to generate matched reference and alternative inputs. For indels, the flanking sequence was adjusted so that both alleles could be represented in the fixed-length peak-centered inputs while preserving the variant within the sequence window. Variant effect values were then calculated for each model output (i.e. cell type) as the difference and log2 fold-change between the alternative and reference predictions. In analyses using a background peak set, peak predictions were additionally used to derive percentile-based scores for variant effects.

#### TF-MINDI analysis

We extracted contribution scores from the DeepCC and DeepSN models for the REF and ALT sequences of all caQTL-ASCA peaks in the corresponding cell types using crested.tl.contribution_scores with the integrated gradients method. Next, we extracted seqlets using TF-MINDI v1.2^38^ (RRID:SCR_027436)(tm.pp.extract_seqlets) separately for CC and SN, combining the high-and low-allele sequences, and retained seqlets with p < 0.01. We then computed seqlet similarity to motifs in a reference motif database using tm.pp.calculate_motif_similarity, which was used for downstream annotation of seqlet clusters with transcription factor family / DNA-binding domain information.

#### scooby models

We trained scooby models for CC and SN separately, on snRNA and snATAC multiome cells with cell type annotation and donor assignment^41^. Data was prepared as described in Hingerl et al.^41^ and sequences were downloaded from the scooby resources (https://zenodo.org/records/14051793). Start coordinates were lifted from hg38 to T2T-CHM13v2.0 and regions were extended back to 196,608 bp. Assignment of sequences to folds was preserved, using fold 3 regions as test sequences and fold 4 regions as validation sequences. Genes and regions overlapping with test and validation sequences were removed as features from the dataset. Additionally, genes and regions appearing in less than 1% of the cells were excluded. Embeddings obtained by Poisson-MultiVI (v1.1.2) were used as input for the scooby models, which were trained with batch size 1 and one H100 GPU. Trained scooby models were used to predict variant effects on gene expression per cell, by summing predicted expression profiles for bins overlapping the gene’s exons. Predictions per cell type were obtained by summing predicted values for each cell of the respective cell type, adding a +1 pseudocount and log2-transforming the values. Variant effect values were then calculated per cell type, as the difference between the cell type prediction (obtained as explained above) of the alternative and reference sequence.

#### Defining GWAS loci

The *pfile* function from PLINK 2.0^83^ (RRID:SCR_001757) was applied to calculate r^2^ values between the 157 leads from recent GWAS study^11^ and 75,193,455 1000G variants from Byrska-Bishop et al.^87^(hg38, PRJEB55077). 93,528 lead-variant pairs were obtained by constraining this function to a 2Mbp window, an r^2^ threshold of 0.05, a minor allele frequency threshold of 0.001, maximal alleles equal to 2 and r^2^-phasing. Afterwards, proximity regions were constructed by finding the furthest variants above an R-squared threshold of 0.6 on either side of each lead variant. This threshold was chosen manually from visualizing different thresholds at each lead variant. The distance to the lead variant - which creates the region - was edited to be 1500 bps minimally on either side making the smallest region size 3,000bp. Six lead variants that were not found in the 1000G set were also assigned this minimal distance. The leads were lifted to T2T-CHM13v2.0 coordinates^68^ using pyLiftover (v0.4.1; RRID:SCR_006646) with the Hg38 to Hs1 USCS chain file^84^ and the left and right distances were kept. Two leads failed liftover resulting in 155 regions which are then used to subset the 53,707,139 donor variants. BEDTools^76^ (2.30.0; RRID:SCR_006646) found 163,695 unique variant hits in the regions that together cover 0.538% of the human genome.

## Availability statement

All data, code, key lab material, and protocols used or created in this study will be, or have already been, deposited in public repositories. These are all listed in the Key Resource Table (https://zenodo.org/records/19132348).

## Data Availability

The WGS data used in this study is stored on EGA under the study accession number EGAS50000000921 and EGAS50000001689. Processed transcriptome single cell count matrices are available via Scope: https://scope.aertslab.org/#/ASAP_brain/*/welcome. Coverage bigwigs for snATAC cell types can be downloaded from https://ucsctracks.aertslab.org/papers/ASAP_paper/. Raw and processed scRNA-seq and scATAC-seq data per donor are already available via 10.5281/zenodo.19034709 and are simultaneously being prepared for release officially via ASAP CRN Cloud (https://cloud.parkinsonsroadmap.org/, https://doi.org/10.5281/zenodo.18988743, https://doi.org/10.5281/zenodo.18988717, https://doi.org/10.5281/zenodo.18988735, https://doi.org/10.5281/zenodo.18988729, https://doi.org/10.5281/zenodo.18988753, https://doi.org/10.5281/zenodo.18988768, https://doi.org/10.5281/zenodo.18988761). CREsted models developed in this paper are available in the CREsted model repository: https://crested.readthedocs.io/en/latest/models/ASAP/index.html.

## Code Availability

The WGS data processing pipeline is available at https://github.com/AlexanRNA/nanowgs/ and is stored on Zenodo 10.5281/zenodo.13385064. The code to reproduce figures can be found in https://github.com/aertslab/ASAP/.

## Acknowledgements

We thank Edinburgh Brain Bank (University of Edinburgh, UK), the Banner Sun Health Research Institute (USA), and the Queen Square Brain Bank for Neurological Disorders (UCL Queen Square Institute of Neurology, UK) for providing samples and their assistance. We would like to thank Chris Flerin and Florian De Rop for their contribution during the initial stages of the project. The computing resources were provided by Flemish Supercomputer Centrum and VIB Data Core. We would like to acknowledge Leuven Genomics Core, VIB Single Cell Core, VIB Nucleomics Core, and VIB Tech Watch for their assistance. We are grateful to Andrew B. Stergachis for providing the Hia5 expression construct and the lab of Zeger Debyser for their assistance with recombinant protein purification. We would like to thank the EMBL Protein Expression and Purification Core Facility for producing recombinant Tn5 protein. We would also like to thank Laura D. Martens and Johannes C. Hingerl for their advice on the scooby models. JD was a postdoctoral fellow of the FWO (Grant number: 12J6921N) and acknowledges support from VIB and KU Leuven Internal Funds (C14/22/125 SymBioSys). AP is a PhD fellow supported by KU Leuven (DB/23/007/bm) and the FWO (Grant number: 1159725N), OS is a Postdoctoral fellow of the FWO (Grant number: 12AZN24N). This research was funded by Aligning Science Across Parkinson’s (ASAP-000430 and ASAP-025179) through the Michael J. Fox Foundation for Parkinson’s Research (MJFF) and CZI (DI2-0000000068). We are grateful to the Banner Sun Health Research Institute Brain and Body Donation Program of Sun City, Arizona for the provision of human biological materials. The Brain and Body Donation Program has been supported by the National Institute of Neurological Disorders and Stroke (U24 NS072026 National Brain and Tissue Resource for Parkinson’s Disease and Related Disorders), the National Institute on Aging (P30 AG019610 and P30AG072980, Arizona Alzheimer’s Disease Center), the Arizona Department of Health Services (contract 211002, Arizona Alzheimer’s Research Center), the Arizona Biomedical Research Commission (contracts 4001, 0011, 05-901 and 1001 to the Arizona Parkinson’s Disease Consortium) and the Michael J. Fox Foundation for Parkinson’s Research. Elements of Fig 1A were obtained from BioRender.

## Author contributions

Conceptualization: SA, JD

Computational analysis: OS, AP, JDM, VK, GH, BS, ADB, LM, KD Subject neuropathological characterization: ACH, BTG, SGE

Data collection, processing, and curation: KT, AM, GH, AP, JDM, OS, JG Experiments and sample preparation: KT, AM

Resources: SA, TV, KV, SS, SM

Software implementation and testing: OS, AP, JDM, VK, GH, JD, BS Visualization: OS, AP, JDM, BS

Writing – original draft: SA, OS, AP, JDM, KT, VK, BS

Writing – review & editing: SA, OS, AP, JDM, KT, VK, JD, TV, ACH, BTG, SGE

## Declaration on competing interests

The authors declare no competing interests.

## Notes

### Competing Interest Statement

The authors have declared no competing interest.

### Summary of Updates

- added supplementary tables - added key resource table and path to CREsted models - minor changes in the manuscript text

## References

1. Brown, C. D., Mangravite, L. M. & Engelhardt, B. E. Integrative Modeling of eQTLs and Cis-Regulatory Elements Suggests Mechanisms Underlying Cell Type Specificity of eQTLs. PLoS Genet. 9, e1003649 (2013).

2. LifeLines Cohort Study et al. Single-cell RNA sequencing identifies celltype-specific cis-eQTLs and co-expression QTLs. Nat. Genet. 50, 493–497 (2018).

3. Pampari, A. et al. ChromBPNet: bias factorized, base-resolution deep learning models of chromatin accessibility reveal cis-regulatory sequence syntax, transcription factor footprints and regulatory variants. Preprint at 10.1101/2024.12.25.630221 (2024).

4. Kempynck, N. et al. CREsted: modeling genomic and synthetic cell type-specific enhancers across tissues and species. Preprint at 10.1101/2025.04.02.646812 (2025).

5. De Winter, S., Konstantakos, V. & Aerts, S. Modelling and design of transcriptional enhancers. Nat. Rev. Bioeng. 3, 374–389 (2025).

6. Avsec, Ž., et al. Advancing regulatory variant effect prediction with AlphaGenome. Nature 649, 1206–1218 (2026).

7. Johansen, N. J. et al. Cross-species consensus atlas of the primate basal ganglia. Preprint at 10.64898/2025.12.15.694496 (2025).

8. Siletti, K. et al. Transcriptomic diversity of cell types across the adult human brain. Science 382, eadd7046 (2023).

9. Mannens, C. C. A. et al. Chromatin accessibility during human first-trimester neurodevelopment. Nature 647, 179–186 (2025).

10. Nalls, M. A. et al. Identification of novel risk loci, causal insights, and heritable risk for Parkinson’s disease: a meta-analysis of genome-wide association studies. Lancet Neurol. 18, 1091–1102 (2019).

11. The Global Parkinson’s Genetics Program (GP2) & Leonard, H. L. Novel Parkinson’s Disease Genetic Risk Factors Within and Across European Populations. Preprint at 10.1101/2025.03.14.24319455 (2025).

12. Blauwendraat, C., Nalls, M. A. & Singleton, A. B. The genetic architecture of Parkinson’s disease. Lancet Neurol. 19, 170–178 (2020).

13. Soldner, F. et al. Parkinson-associated risk variant in distal enhancer of α-synuclein modulates target gene expression. Nature 533, 95–99 (2016).

14. Langston, R. G. et al. Association of a common genetic variant with Parkinson’s disease is mediated by microglia. Sci. Transl. Med. 14, eabp8869 (2022).

15. Prahl, J. D., Pierce, S. E., Van Der Schans, E. J. C., Coetzee, G. A. & Tyson, T. The Parkinson’s disease variant rs356182 regulates neuronal differentiation independently from alpha-synuclein. Hum. Mol. Genet. 32, 1–14 (2023).

16. Corces, M. R. et al. Single-cell epigenomic analyses implicate candidate causal variants at inherited risk loci for Alzheimer’s and Parkinson’s diseases. Nat. Genet. 52, 1158–1168 (2020).

17. Álvarez Jerez, P., et al. African ancestry neurodegeneration risk variant disrupts an intronic branchpoint in GBA1. Nat. Struct. Mol. Biol. 31, 1955–1963 (2024).

18. Braak, H. et al. Staging of brain pathology related to sporadic Parkinson’s disease. Neurobiol. Aging 24, 197–211 (2003).

19. Stergachis, A. B., Debo, B. M., Haugen, E., Churchman, L. S. & Stamatoyannopoulos, J. A. Single-molecule regulatory architectures captured by chromatin fiber sequencing. Science 368, 1449–1454 (2020).

20. Kolmogorov, M. et al. Scalable Nanopore sequencing of human genomes provides a comprehensive view of haplotype-resolved variation and methylation. Nat. Methods 20, 1483–1492 (2023).

21. Xu, C. et al. Probabilistic harmonization and annotation of single-cell transcriptomics data with deep generative models. Mol. Syst. Biol. 17, e9620 (2021).

22. Bravo González-Blas, C., et al. SCENIC+: single-cell multiomic inference of enhancers and gene regulatory networks. Nat. Methods 20, 1355–1367 (2023).

23. Hoffman, G. E. et al. Efficient differential expression analysis of large-scale single cell transcriptomics data using dreamlet. Preprint at 10.1101/2023.03.17.533005 (2023).

24. Hill, M. S., Vande Zande, P. & Wittkopp, P. J. Molecular and evolutionary processes generating variation in gene expression. Nat. Rev. Genet. 22, 203–215 (2021).

25. Atak, Z. K. et al. Interpretation of allele-specific chromatin accessibility using cell state–aware deep learning. Genome Res. (2021) doi:10.1101/gr.260851.120.

26. Taylor-Weiner, A. et al. Scaling computational genomics to millions of individuals with GPUs. Genome Biol. 20, 228 (2019).

27. van de Geijn, B., McVicker, G., Gilad, Y. & Pritchard, J. K. WASP: allele-specific software for robust molecular quantitative trait locus discovery. Nat. Methods 12, 1061–1063 (2015).

28. Cuomo, A. S. E. et al. Impact of rare and common genetic variation on cell type-specific gene expression in human blood. Preprint at 10.1101/2025.03.20.25324352 (2025).

29. Kanai, M. et al. Population-scale multiome immune cell atlas reveals complex disease drivers. Preprint at 10.1101/2025.11.25.25340489 (2025).

30. Benaglio, P. et al. Mapping genetic effects on cell type-specific chromatin accessibility and annotating complex immune trait variants using single nucleus ATAC-seq in peripheral blood. PLoS Genet. 19, e1010759 (2023).

31. Natri, H. M. et al. Cell-type-specific and disease-associated expression quantitative trait loci in the human lung. Nat. Genet. 56, 595–604 (2024).

32. Xue, A. et al. Genetic regulation of cell type–specific chromatin accessibility shapes immune function and disease risk. Preprint at 10.1101/2025.08.27.25334533 (2025).

33. Hecker, N. et al. Enhancer-driven cell type comparison reveals similarities between the mammalian and bird pallium. Science 387, eadp3957 (2025).

34. Li, Y. E. et al. A comparative atlas of single-cell chromatin accessibility in the human brain. Science 382, eadf7044 (2023).

35. Zemke, N. R. et al. Conserved and divergent gene regulatory programs of the mammalian neocortex. Nature 624, 390–402 (2023).

36. Deplancke, B., Alpern, D. & Gardeux, V. The Genetics of Transcription Factor DNA Binding Variation. Cell 166, 538–554 (2016).

37. Abramov, S. et al. Landscape of allele-specific transcription factor binding in the human genome. Nat. Commun. 12, 2751 (2021).

38. De Winter, S. et al. System-wide extraction of cis-regulatory rules from sequence-to-function models in human neural development. Preprint at 10.64898/2026.01.14.699402 (2026).

39. Schreiber, J. tangermeme: A toolkit for understanding cis-regulatory logic using deep learning models. Preprint at 10.1101/2025.08.08.669296 (2025).

40. Linder, J., Srivastava, D., Yuan, H., Agarwal, V. & Kelley, D. R. Predicting RNA-seq coverage from DNA sequence as a unifying model of gene regulation. Nat. Genet. 57, 949–961 (2025).

41. Hingerl, J. C. et al. scooby: modeling multimodal genomic profiles from DNA sequence at single-cell resolution. Nat. Methods 22, 2275–2285 (2025).

42. Martens, L. D., Fischer, D. S., Yépez, V. A., Theis, F. J. & Gagneur, J. Modeling fragment counts improves single-cell ATAC-seq analysis. Nat. Methods 21, 28–31 (2024).

43. Hamza, T. H. et al. Common genetic variation in the HLA region is associated with late-onset sporadic Parkinson’s disease. Nat. Genet. 42, 781–785 (2010).

44. Ahmed, I. et al. Association between Parkinson’s disease and the *HLA-DRB1* locus. Mov. Disord. 27, 1104–1110 (2012).

45. Yu, E. et al. Fine mapping of the HLA locus in Parkinson’s disease in Europeans. Npj Park. Dis. 7, 84 (2021).

46. Zody, M. C. et al. Evolutionary toggling of the MAPT 17q21.31 inversion region. Nat. Genet. 40, 1076–1083 (2008).

47. Smaili, I. et al. A Specific Diplotype H1j/H2 of the MAPT Gene Could Be Responsible for Parkinson’s Disease with Dementia. Case Rep. Genet. 2020, 1–5 (2020).

48. Pedicone, C., Weitzman, S. A., Renton, A. E. & Goate, A. M. Unraveling the complex role of MAPT-containing H1 and H2 haplotypes in neurodegenerative diseases. Mol. Neurodegener. 19, 43 (2024).

49. Bayle, E. D. et al. Carboxylesterase Notum Is a Druggable Target to Modulate Wnt Signaling. J. Med. Chem. 64, 4289–4311 (2021).

50. Teranishi, Y. et al. Proton myo-inositol cotransporter is a novel γ-secretase associated protein that regulates Aβ production without affecting Notch cleavage. FEBS J. 282, 3438–3451 (2015).

51. Hormozdiari, F. et al. Colocalization of GWAS and eQTL Signals Detects Target Genes. Am. J. Hum. Genet. 99, 1245–1260 (2016).

52. The GTEx Consortium et al. The GTEx Consortium atlas of genetic regulatory effects across human tissues. Science 369, 1318–1330 (2020).

53. Connally, N. J. et al. The missing link between genetic association and regulatory function. eLife 11, e74970 (2022).

54. Mostafavi, H., Spence, J. P., Naqvi, S. & Pritchard, J. K. Systematic differences in discovery of genetic effects on gene expression and complex traits. Nat. Genet. 55, 1866–1875 (2023).

55. Uhlén, M. et al. Tissue-based map of the human proteome. Science 347, 1260419 (2015).

56. Shen, Y. et al. BST1 rs4698412 allelic variant increases the risk of gait or balance deficits in patients with Parkinson’s disease. CNS Neurosci. Ther. 25, 422–429 (2019).

57. Guerreiro, S., Privat, A.-L., Bressac, L. & Toulorge, D. CD38 in Neurodegeneration and Neuroinflammation. Cells 9, 471 (2020).

58. Xu, H.-L. et al. The impact of BST1 rs4698412 variant on Parkinson’s disease progression in a longitudinal study. Front. Aging Neurosci. 17, 1570347 (2025).

59. Basak, S. et al. Novel Roles for the Ectoenzyme CD38 in the Maintenance of Transcriptional and Metabolic Homeostasis in Astrocytes. Glia 74, e70112 (2026).

60. Chang, D. et al. A meta-analysis of genome-wide association studies identifies 17 new Parkinson’s disease risk loci. Nat. Genet. 49, 1511–1516 (2017).

61. Huang, C. et al. Personal transcriptome variation is poorly explained by current genomic deep learning models. Nat. Genet. 55, 2056–2059 (2023).

62. Sasse, A. et al. Benchmarking of deep neural networks for predicting personal gene expression from DNA sequence highlights shortcomings. Nat. Genet. 55, 2060–2064 (2023).

63. Zhou, W. et al. Efficient and accurate mixed model association tool for single-cell eQTL analysis. Preprint at 10.1101/2024.05.15.24307317 (2024).

64. Beach, T. G. et al. Arizona Study of Aging and Neurodegenerative Disorders and Brain and Body Donation Program. Neuropathol. Off. J. Jpn. Soc. Neuropathol. 35, 354–389 (2015).

65. Li, H. Minimap2: pairwise alignment for nucleotide sequences. Bioinformatics 34, 3094–3100 (2018).

66. Zheng, Z. et al. Symphonizing pileup and full-alignment for deep learning-based long-read variant calling. Nat. Comput. Sci. 2, 797–803 (2022).

67. Danecek, P. et al. Twelve years of SAMtools and BCFtools. GigaScience 10, giab008 (2021).

68. Nurk, S. et al. The complete sequence of a human genome. Science 376, 44–53 (2022).

69. De Coster, W. & Rademakers, R. NanoPack2: population-scale evaluation of long-read sequencing data. Bioinformatics 39, btad311 (2023).

70. Pedersen, B. S. & Quinlan, A. R. Mosdepth: quick coverage calculation for genomes and exomes. Bioinformatics 34, 867–868 (2018).

71. Smolka, M. et al. Detection of mosaic and population-level structural variants with Sniffles2. Nat. Biotechnol. 42, 1571–1580 (2024).

72. Lin, J.-H., Chen, L.-C., Yu, S.-C. & Huang, Y.-T. LongPhase: an ultra-fast chromosome-scale phasing algorithm for small and large variants. Bioinformatics 38, 1816–1822 (2022).

73. Cheetham, S. W., Kindlova, M. & Ewing, A. D. Methylartist: tools for visualizing modified bases from nanopore sequence data. Bioinformatics 38, 3109–3112 (2022).

74. Pedersen, B. S. et al. Somalier: rapid relatedness estimation for cancer and germline studies using efficient genome sketches. Genome Med. 12, 62 (2020).

75. Heumos, L. et al. Best practices for single-cell analysis across modalities. Nat. Rev. Genet. 24, 550–572 (2023).

76. Montserrat-Ayuso, T. & Esteve-Codina, A. High content of nuclei-free low-quality cells in reference single-cell atlases: a call for more stringent quality control using nuclear fraction. BMC Genomics 25, 1124 (2024).

77. Bakken, T. E. et al. Comparative cellular analysis of motor cortex in human, marmoset and mouse. Nature 598, 111–119 (2021).

78. Kamath, T. et al. Single-cell genomic profiling of human dopamine neurons identifies a population that selectively degenerates in Parkinson’s disease. Nat. Neurosci. 25, 588–595 (2022).

79. Lopez, R., Regier, J., Cole, M. B., Jordan, M. I. & Yosef, N. Deep generative modeling for single-cell transcriptomics. Nat. Methods 15, 1053–1058 (2018).

80. Casper, J. et al. The UCSC Genome Browser database: 2026 update. Nucleic Acids Res. 54, D1331–D1335 (2026).

81. Huang, X. & Huang, Y. Cellsnp-lite: an efficient tool for genotyping single cells. Bioinformatics 37, 4569–4571 (2021).

82. Huang, Y., McCarthy, D. J. & Stegle, O. Vireo: Bayesian demultiplexing of pooled single-cell RNA-seq data without genotype reference. Genome Biol. 20, 273 (2019).

83. Chang, C. C. et al. Second-generation PLINK: rising to the challenge of larger and richer datasets. Gigascience 4, s13742-015-0047–8 (2015).

84. Heumos, L. et al. Pertpy: an end-to-end framework for perturbation analysis. Nat. Methods 23, 350–359 (2026).

85. Sigalova, O. M. et al. Integrating genetic variation with deep learning provides context for variants impacting transcription factor binding during embryogenesis. Genome Res. 35, 1138–1153 (2025).

86. Benjamini, Y. & Hochberg, Y. Controlling the False Discovery Rate: A Practical and Powerful Approach to Multiple Testing. J. R. Stat. Soc. Ser. B Methodol. 57, 289–300 (1995).

87. Byrska-Bishop, M. et al. High-coverage whole-genome sequencing of the expanded 1000 Genomes Project cohort including 602 trios. Cell 185, 3426–3440.e19 (2022).

88. Perez, G. et al. The UCSC Genome Browser database: 2025 update. Nucleic Acids Res. 53, D1243–D1249 (2025).

89. Quinlan, A. R. & Hall, I. M. BEDTools: a flexible suite of utilities for comparing genomic features. Bioinformatics 26, 841–842 (2010).

