## Supplementary figures for "Modeling *cis*-regulatory variation in human brain enhancers across a large Parkinson’s Disease cohort"

Affiliations
1 Laboratory of Computational Biology, VIB Center for AI & Computational Biology (VIB.AI), Leuven, Belgium
2 VIB-KU Leuven Center for Neuroscience, Leuven, Belgium
3 Department of Human Genetics, KU Leuven, Leuven, Belgium
4 Laboratory of Integrative Cancer Genomics, Department of Oncology, KU Leuven, Leuven, Belgium
5 VIB Center for Cancer Biology, VIB, Leuven, Belgium
6 Aligning Science Across Parkinson's (ASAP) Collaborative Research Network, Chevy Chase, MD 20815, United States
7 KU Leuven Institute for Single Cell Omics (LISCO), KU Leuven, Leuven, Belgium
8 Laboratory of Reproductive Genomics, Department of Human Genetics, KU Leuven, Leuven, Belgium
9 Department of Neurology, Mayo Clinic School of Medicine, Mayo Clinic Arizona, Scottsdale
10 Banner Sun Health Research Institute, Sun City, AZ, United States
11 VIB Center for AI & Computational Biology (VIB.AI), Leuven, Belgium

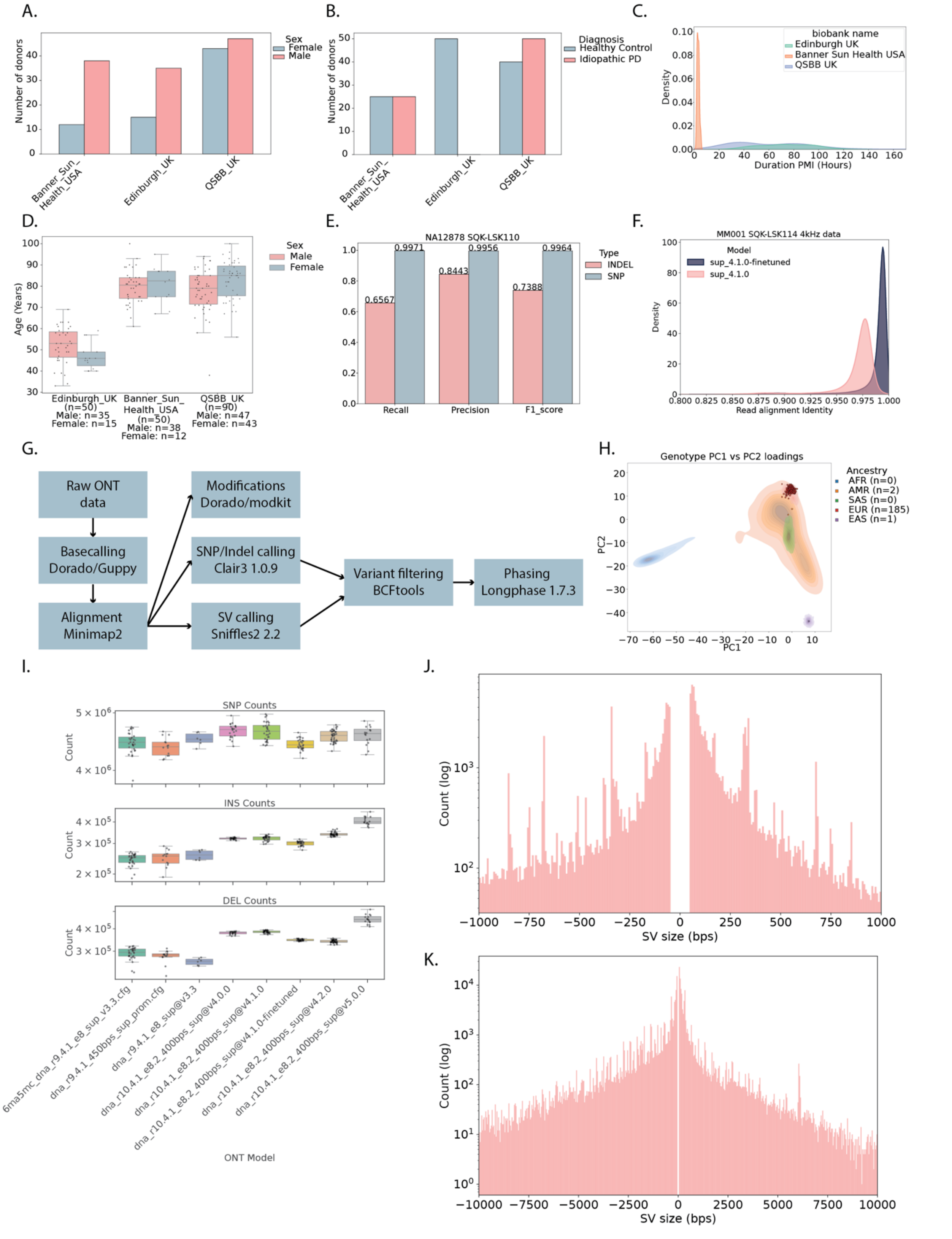


**Supplementary Figure 1: Study cohort and WGS overview and metrics**

**A**. Number of donors per biobank stratified by sex. **B**. Number of donors per biobank stratified by diagnosis. **C**. Postmortem interval distribution in hours colored by biobank. **D**. Age distribution divided by biobank and sex. **E**. Precision, recall and F1 metrics for GIAB SNP and indel truth set for ONT Fiber-seq SQK-LSK100 NA128778 basecalled with a fine-tuned model. **F**. Density distribution of read alignment identity for MM001 SQK-LSK114 ONT Fiber-seq data colored by basecalling model used (dna_r10.4.1_e8.2_400bps_sup@4.1.0 or dna_r10.4.1_e8.2_400bps_sup@4.1.0-finetuned, sup_4.1.0 and sup_4.1.0-finetuned in the legend, respectively). **G**. NanoWGS pipeline overview. **H**. PC1 vs PC2 genotype loadings. Kernel density color represents ancestry distributions while dots represent individual donors in our cohort. **I**. SNP and indel (< 50 bps) counts stratified by basecalling model used. **J,K**. Histogram of SV counts by size for SVs < 1 kbps (J) and SVs <10 kbps (K).

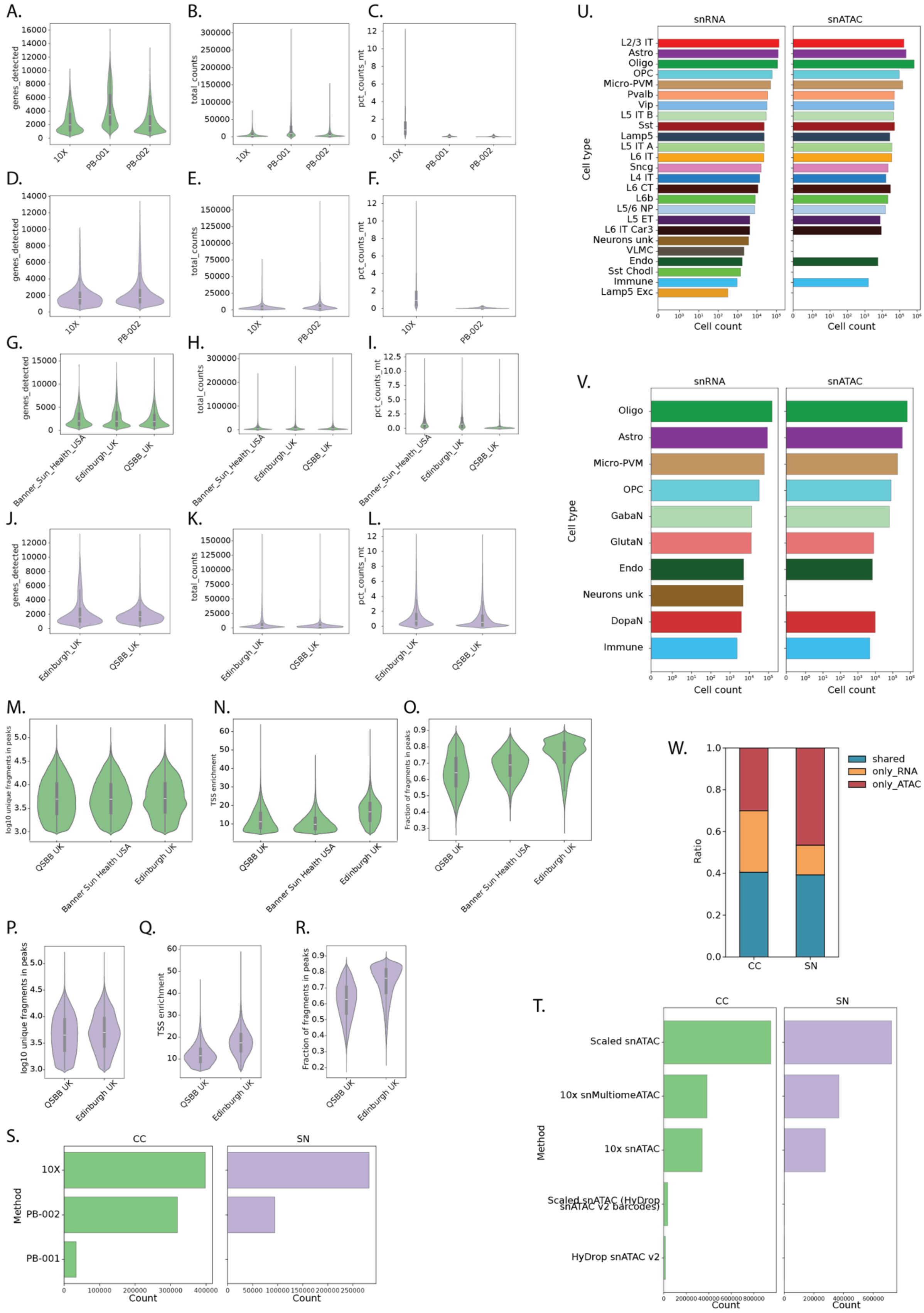


**Supplementary Figure 2: Single-cell multiome atlas quality metrics**

**A-C.** Number of genes detected (A), total UMI counts (B), and % mitochondrial counts (C) in cingulate cortex snRNA stratified by method used (10X, ParseBio 100k kit – PB-001, ParseBio

1M kit – PB-002). **D-F.** Number of genes detected (D), total UMI counts (E), and % mitochondrial counts (F) in substantia nigra snRNA stratified by method used (10X, ParseBio 1M kit – PB-002). **G-I.** Same as (A-C; i.e. cingulate cortex) but stratified by biobank. **J-L.** Same as (D-F; i.e. substantia nigra) but stratified by biobank. **M-O.** Log10 unique fragments in peaks (M), TSS enrichment (N), and fraction of fragments in peaks (O) in cingulate cortex snATAC stratified by biobank. **P-R.** Same as (M-O) for substantia nigra. **S.** Number of transcriptome-called nuclei per method and brain region **T.** Number of chromatin accessibility-called nuclei per method and brain region. **U,V.** Cell counts in cingulate cortex (U) and substantia nigra (V) by cell type for snRNA and snATAC. **W.** Fraction of cell barcodes shared, unique to ATAC and unique to RNA for 10X Multiome nuclei.

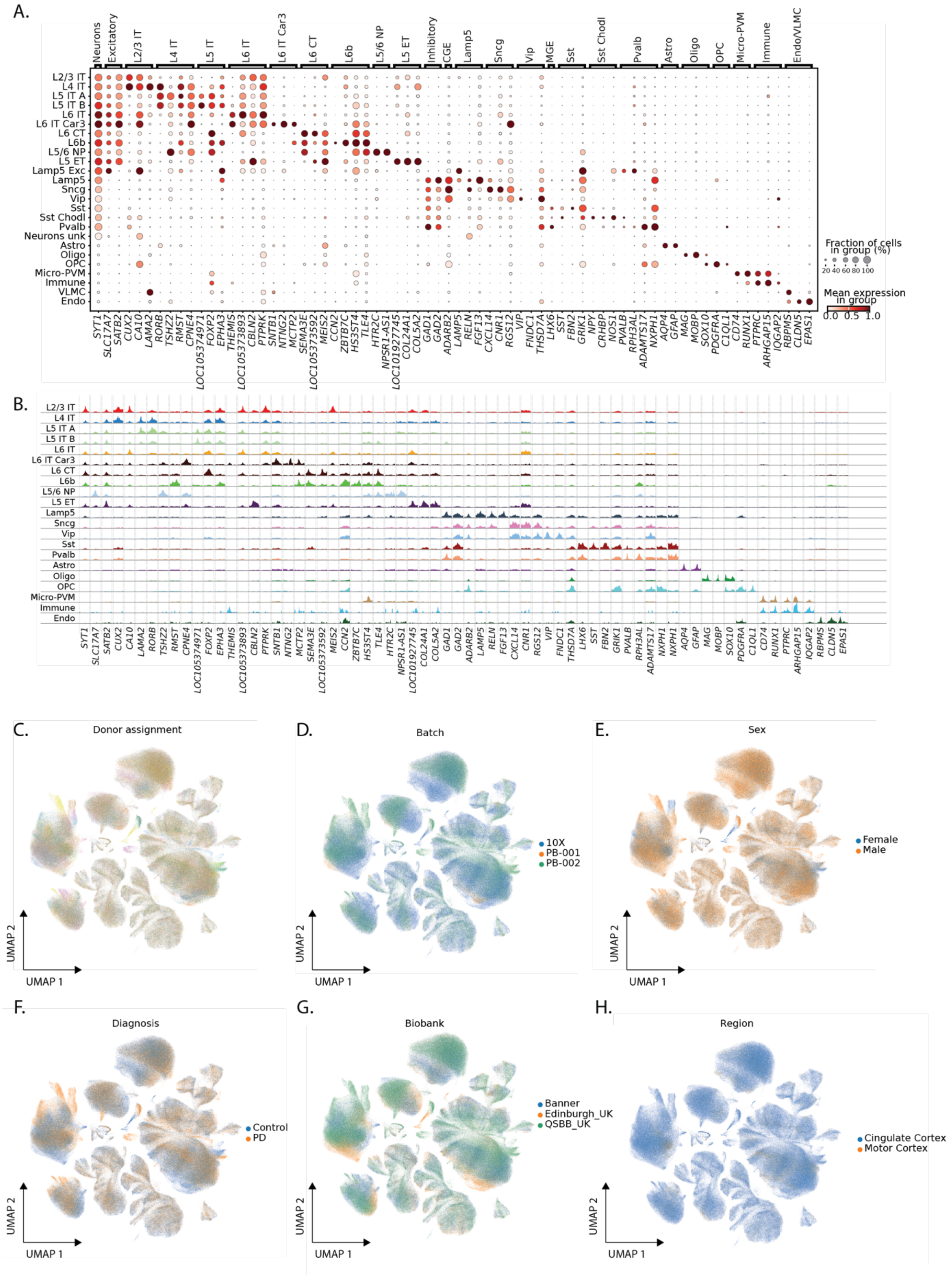


**Supplementary figure 3: Cingulate cortex multiomic atlas overview**

**A**. Average gene expression of select marker genes in cingulate cortex cell types. **B**. Pseudobulk chromatin accessibility profiles in proximity of marker genes in (A). **C-H**. UMAP of cingulate cortex colored by donor assignment (C), batch (D), sex (E), diagnosis (F), biobank (G), and region (H).

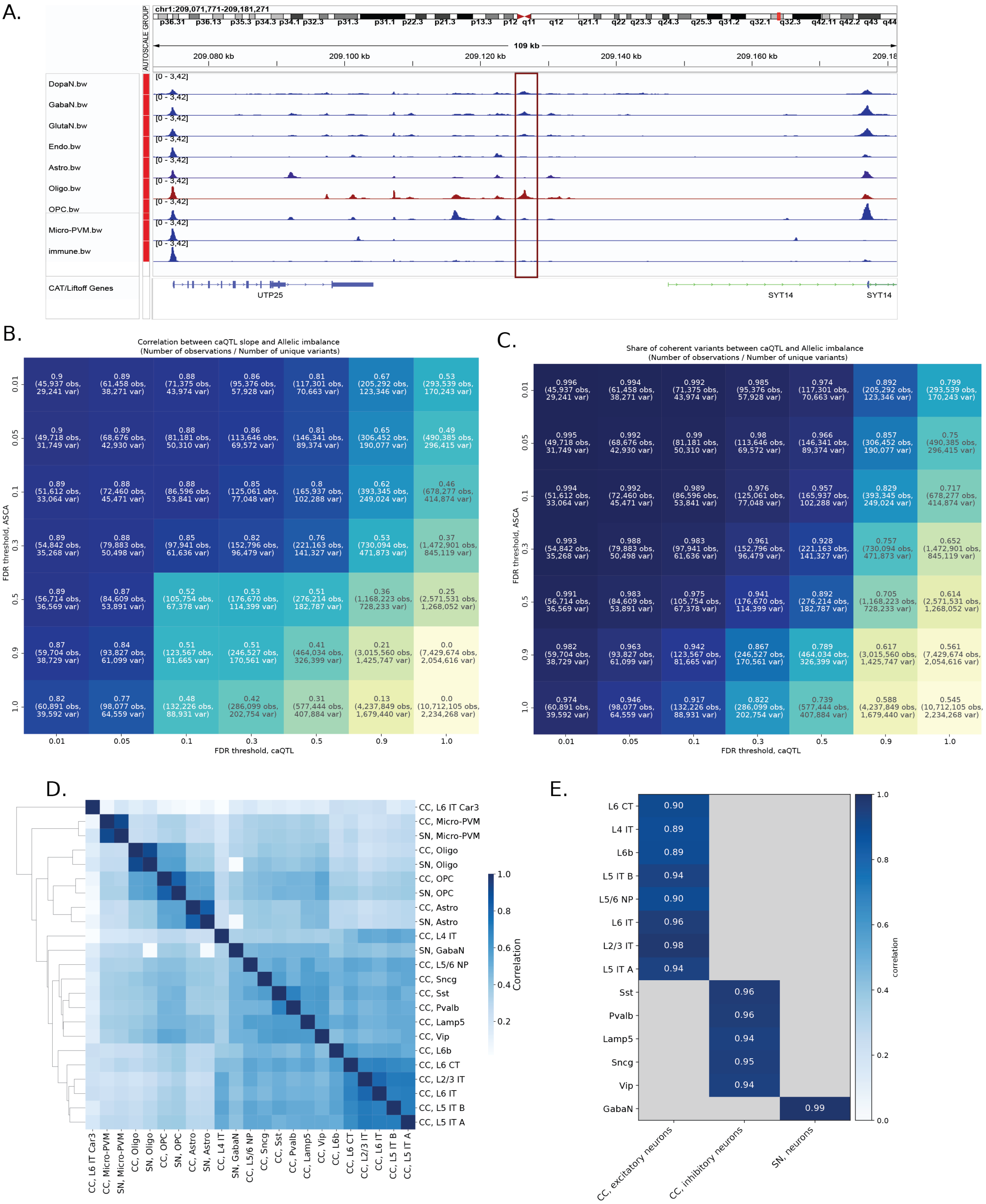


**Supplementary figure 4: Cell type-specific ASCA-caQTL in brain data**

**A**. IGV view of cell type-specific chromatin accessibility bigwig tracks spanning chr1:209,071,771-209,181,271 highlighting an Oligodendrocyte-specific peak in SN. **B**. Correlation between caQTL slope and allelic imbalance at various FDR thresholds for caQTL and ASCA testing. Numbers of observations and unique variants are in brackets. **C**. Fraction of coherent variants between caQTL slope and allelic imbalance at various FDR thresholds for caQTL and ASCA testing. Numbers of observations and unique variants are in brackets. **D**. Correlation of caQTL slopes across cell types. All caQTL & ASCA variants in at least one cell type are considered, caQTL slopes are compared across all cell types where a variant is quantified (regardless of significance). **E**. Correlation between caQTL slopes between broad and granular neuronal cell types for caQTL-ASCA callset with FDR < 0.1 in broad neuronal cell types.

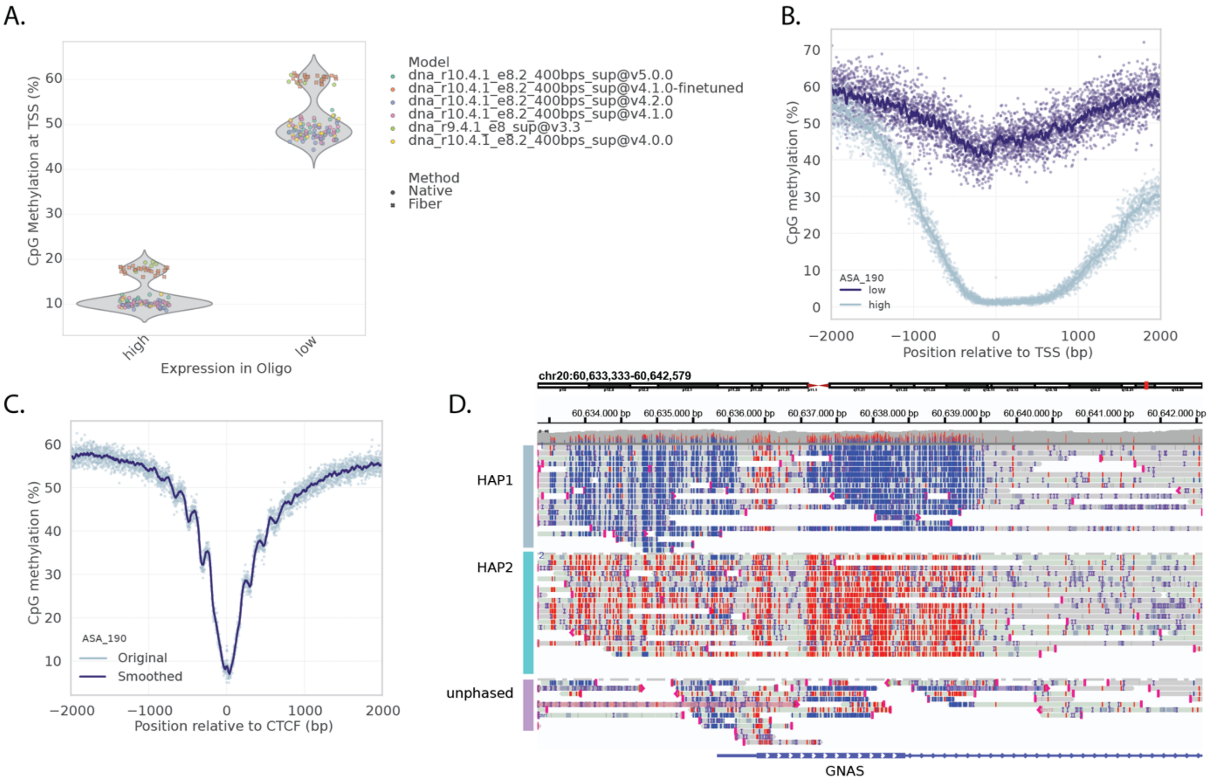


**Supplementary figure 5: Assessment of CpG Methylation in brain LR-WGS data**

**A.** Average CpG methylation in a 4 kb window around TSS for the 10% highest and lowest expressed genes, respectively, in CC Oligodendrocytes. Note that modification calling on older R9.4.1 Nanopore chemistry (SQK-LSK10) and newer R10.4.1 (SQK-LSK114) chemistry Fiber-Seq samples likely overestimates 5mCpG methylation rates. **B**. Average bulk WGS 5mC CpG methylation for donor ASA_190 in a 4 kb window around TSS for the 10% highest and lowest expressed genes in Oligodendrocytes. Lines represent sliding window averages. **C**. Same as (B) around CTCF binding sites. **D**. IGV view of the imprinted *GNAS* promoter (chr20:60,633,333-60,642,579) in donor ASA_190. Reads are phased and colored by methylation status with 5mCpG and canonical CpG colored red and blue, respectively.

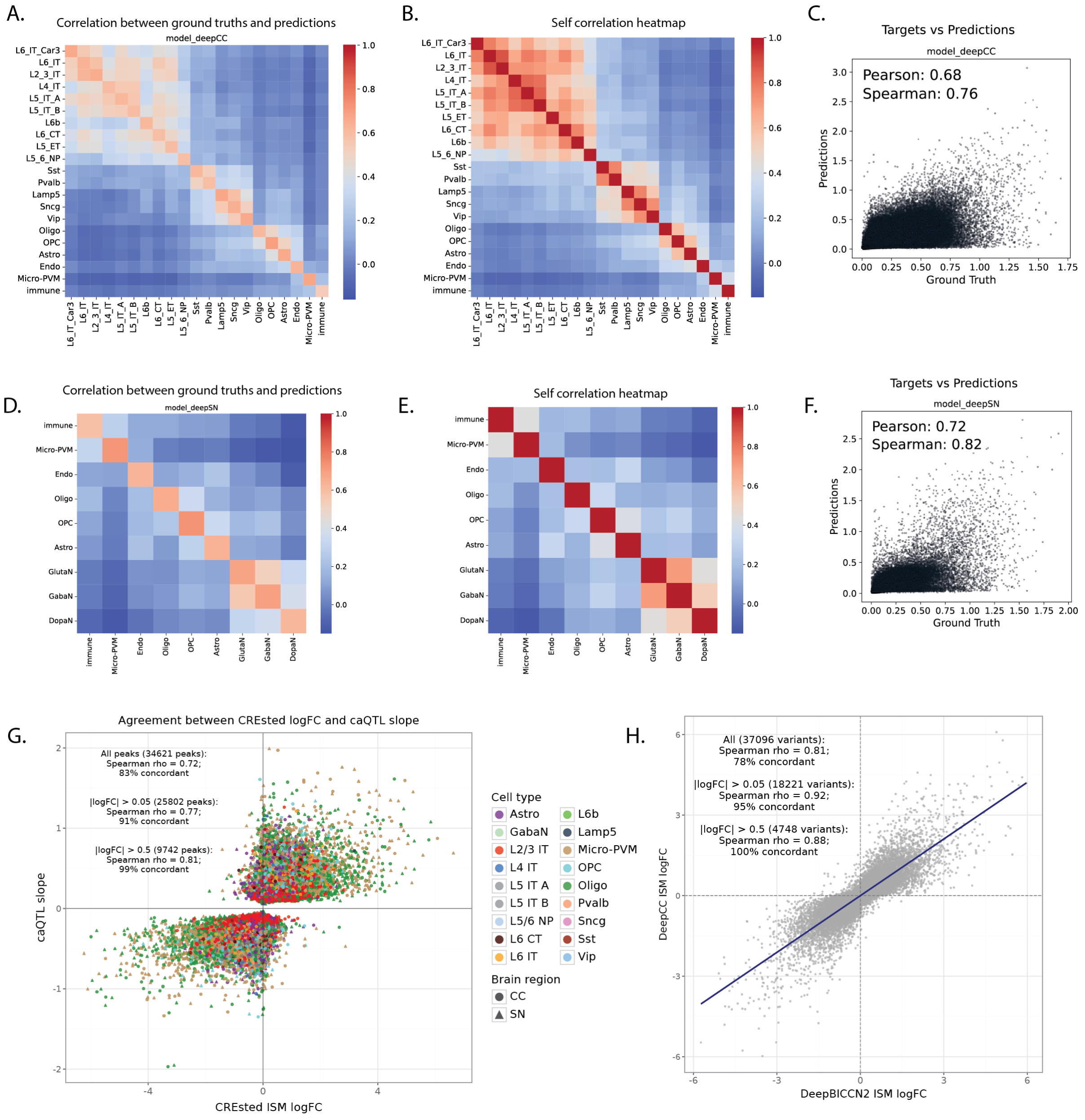


**Supplementary figure 6: Overview of performance of in-house trained CREsted models**
**A.** Correlation between ground truth and predicted chromatin accessibility values for DeepCC model on the test set of highly variable regions (finetuning set). **B.** Self-correlation between ground truth values for DeepCC model. **C.** Ground truth versus predicted chromatin accessibility values on test set from DeepCC model (all cell types). **D.** Correlation between ground truth and predicted chromatin accessibility values for DeepSN model on the test set of highly variable regions (finetuning set). **E.** Self-correlation between ground truth values for deepCC model. **F.** Ground truth versus predicted chromatin accessibility values on test set from DeepSN model (all cell types). **G.** Agreement between CREsted logFC and caQTL slope for all 34,621 caQTL-ASCA peaks. **H.** Agreement between DeepBICCN2 ISM logFC vs DeepCC ISM logFC for all caQTL-ASCA variants tested in both models.

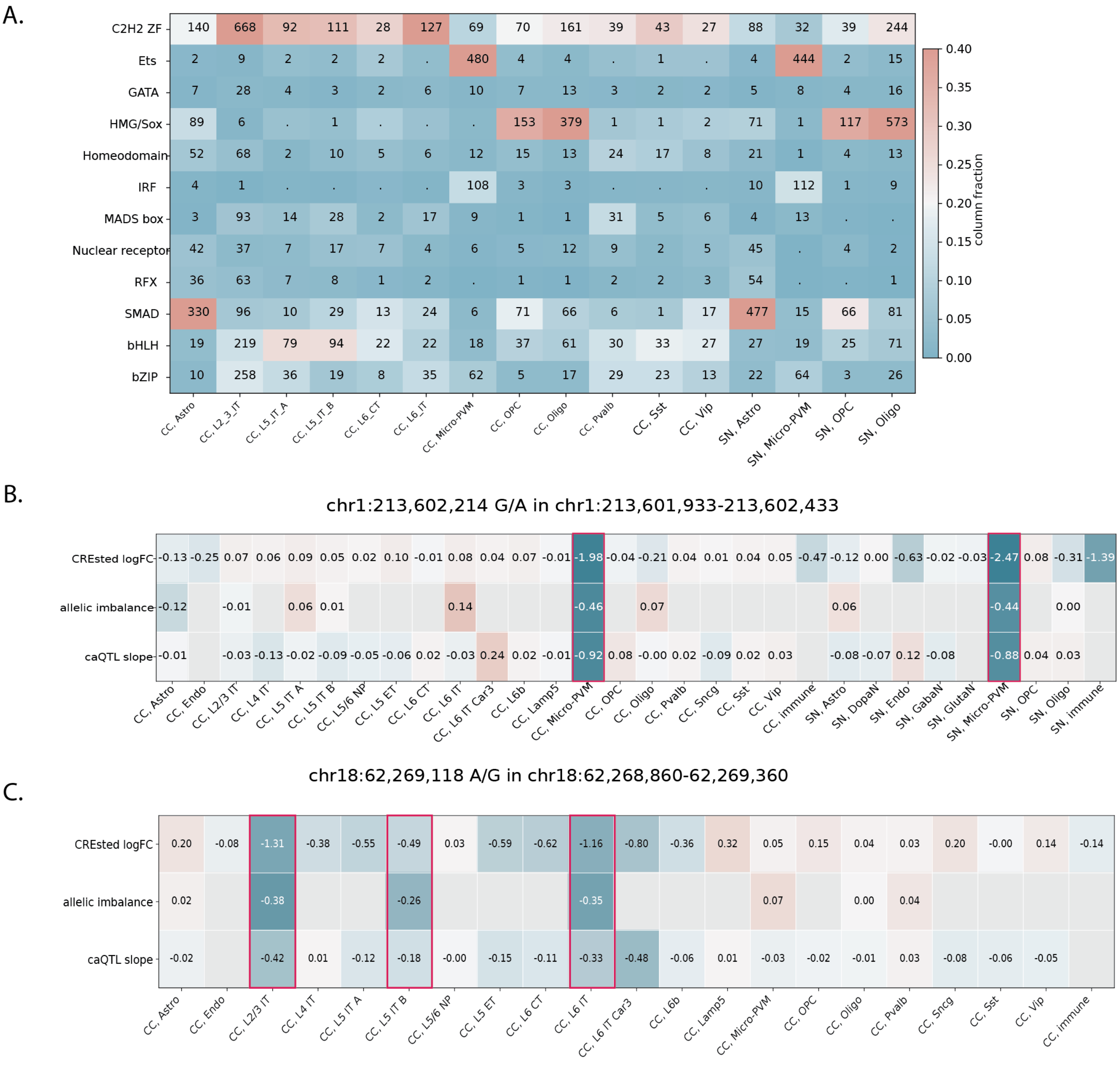

**Supplementary figure 7: Analysis of disrupted cis-regulatory grammar**
**A.** Seqlets affected by caQTL-ASCA variants per cell type. **B**. CREsted logFC, allelic imbalance, and caQTL slope values for all tested cell types across two brain regions for variant chr1:213,602,214 G/A in the ATAC peak chr1:213,601,933-213,602,433. The statistically significant caQTL-ASCA variants are highlighted in red box. **C**. CREsted logFC, allelic imbalance, caQTL slope values for all tested cell types across two brain regions for variant chr18:62,269,118 A/G in the ATAC peak chr18:62,268,860-62,269,360. The statistically significant caQTL-ASCA variants are highlighted in red box.

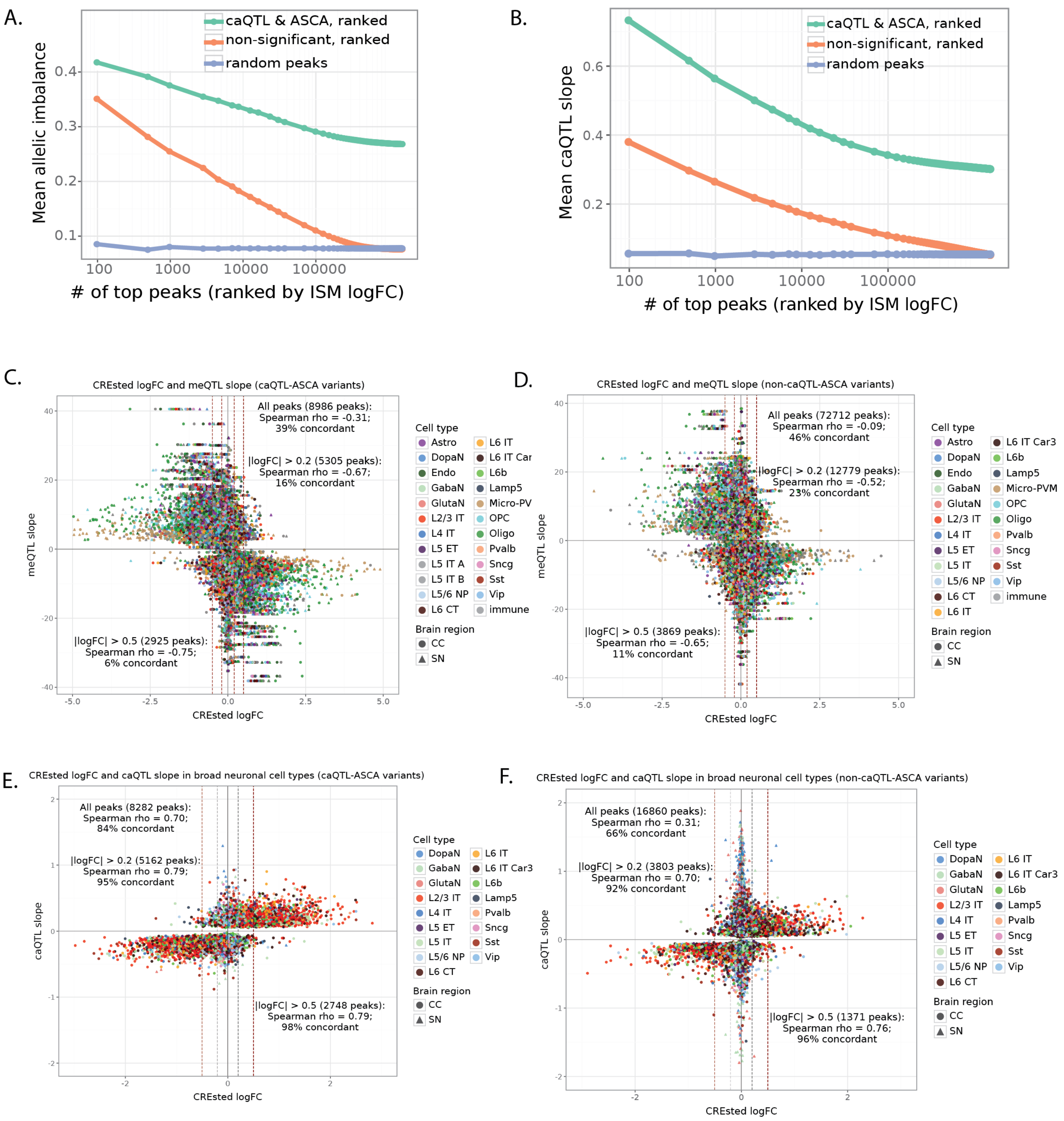


**Supplementary figure 8: CREsted prediction agreement with ASCA/caQTL/meQTL variants**

**A,B.** Rank plot of mean allelic imbalance (A) and mean caQTL slope (B) of peaks ranked by absolute CREsted logFC. Peaks are stratified by whether they contain (non)-significant caQTL-ASCA variants and ranked by CREsted logFC. Values for the same number of randomly selected peaks (not ranked) are shown in blue. **C,D.** CREsted logFC vs meQTL slope for caQTL-ASCA call set variants (C) and for non- caQTL-ASCA variants (D). Only variants with nominal meQTL FDR < 0.1 are shown. **E,F.** CREsted logFC vs caQTL slope in broad cell types for caQTL-ASCA call set variants (E) and for non- caQTL-ASCA variants (F). Only variants with caQTL FDR < 0.1 are shown.

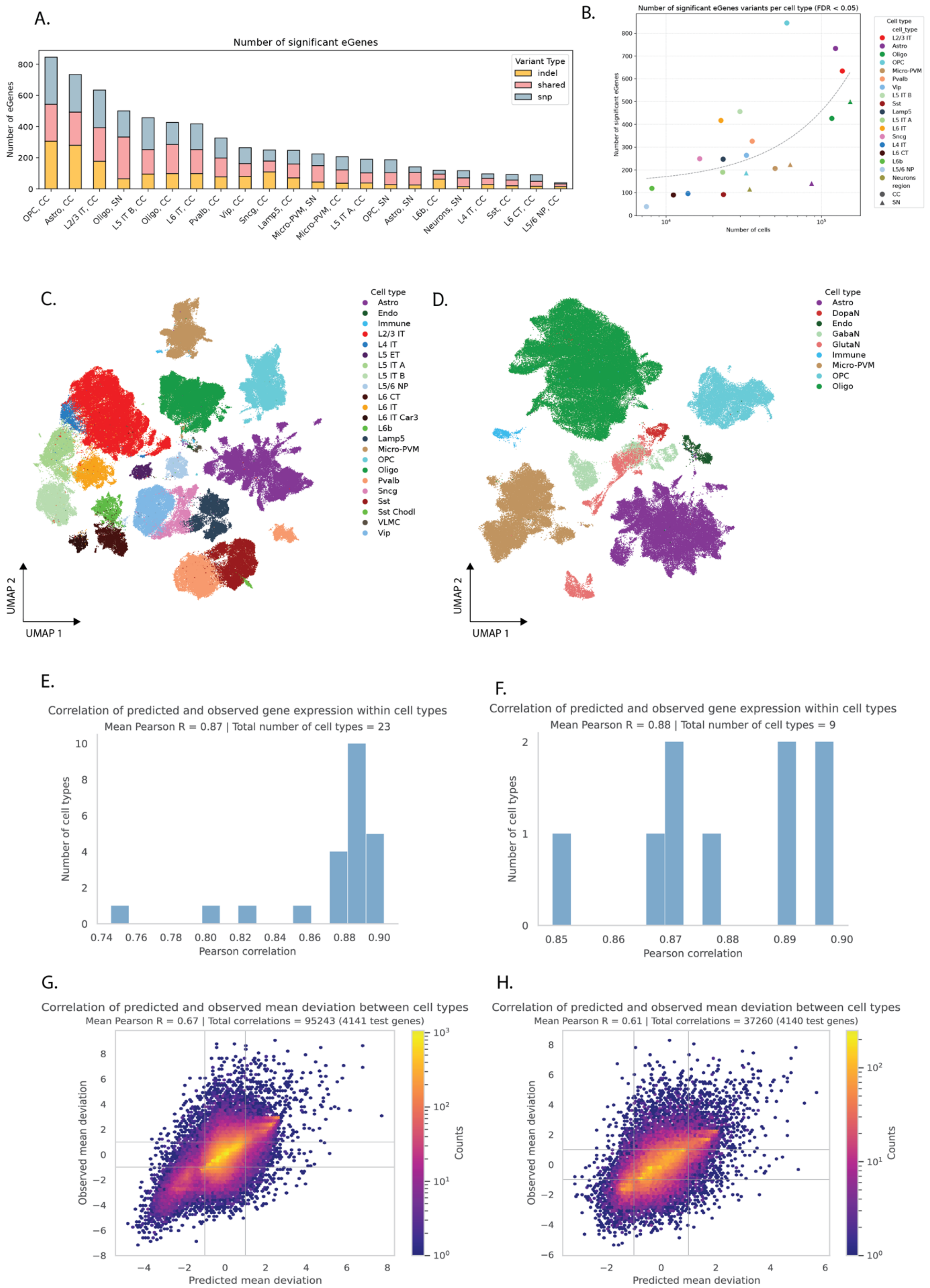


**Supplementary figure 9: eQTL and variant effect prediction with Scooby**

**A.** Bar plot showing the number of significant eGenes (FDR<0.05) for each cell type; colored by variant type (SNP in blue, indel in yellow). ‘Shared’ means the eGene had both a significant indel and SNP. **B**. Number of cells versus the number of significant eGenes per cell type. **C**,**D**. UMAP of common transcriptome and chromatin accessibility space for CC (C) and SN (D). **E,F.** Mean Pearson correlation distribution of scooby predicted and observed gene expression (as counts, summed over the exons) for CC (E) and SN (F) cell types. **G,H.** scooby predicted versus observed gene expression, after removing mean gene expression over all genes per cell-type and removing mean gene expression over all cell types per gene for CC (G) and SN (H). Dots, colored by density, represent predicted values for all genes and all cell types against observed values for all genes and all cell types.

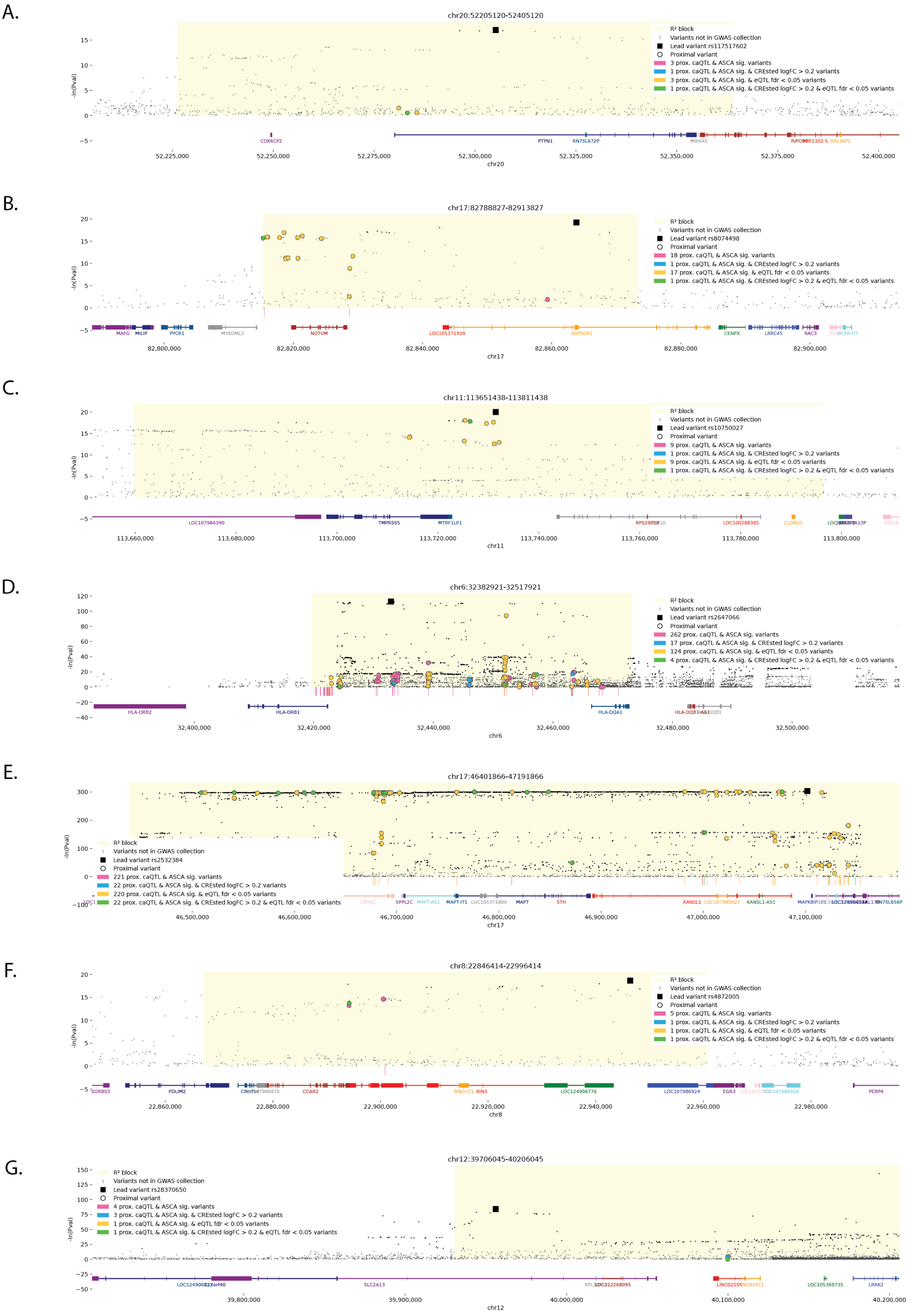


**Supplementary figure 10: Identified cis-regulatory variation at PD GWAS loci**

**A-G.** Manhattan plots of *PTPN1* (A), *NOTUM* (B), *TMPRSS5* (C) *HLA* (D), *MAPT-KANSL1* (E), *BIN3* (F), and *LRRK2* (G) loci with variants as gray dots and significant GWAS hits ($p < 5x10^{-8}$) in black. Identified caQTL/ASCA/CREsted/eQTL variants and GWAS lead variants within $R^2$-identified LD blocks (light yellow) are highlighted and color-coded. Gene annotations are shown below the plots.

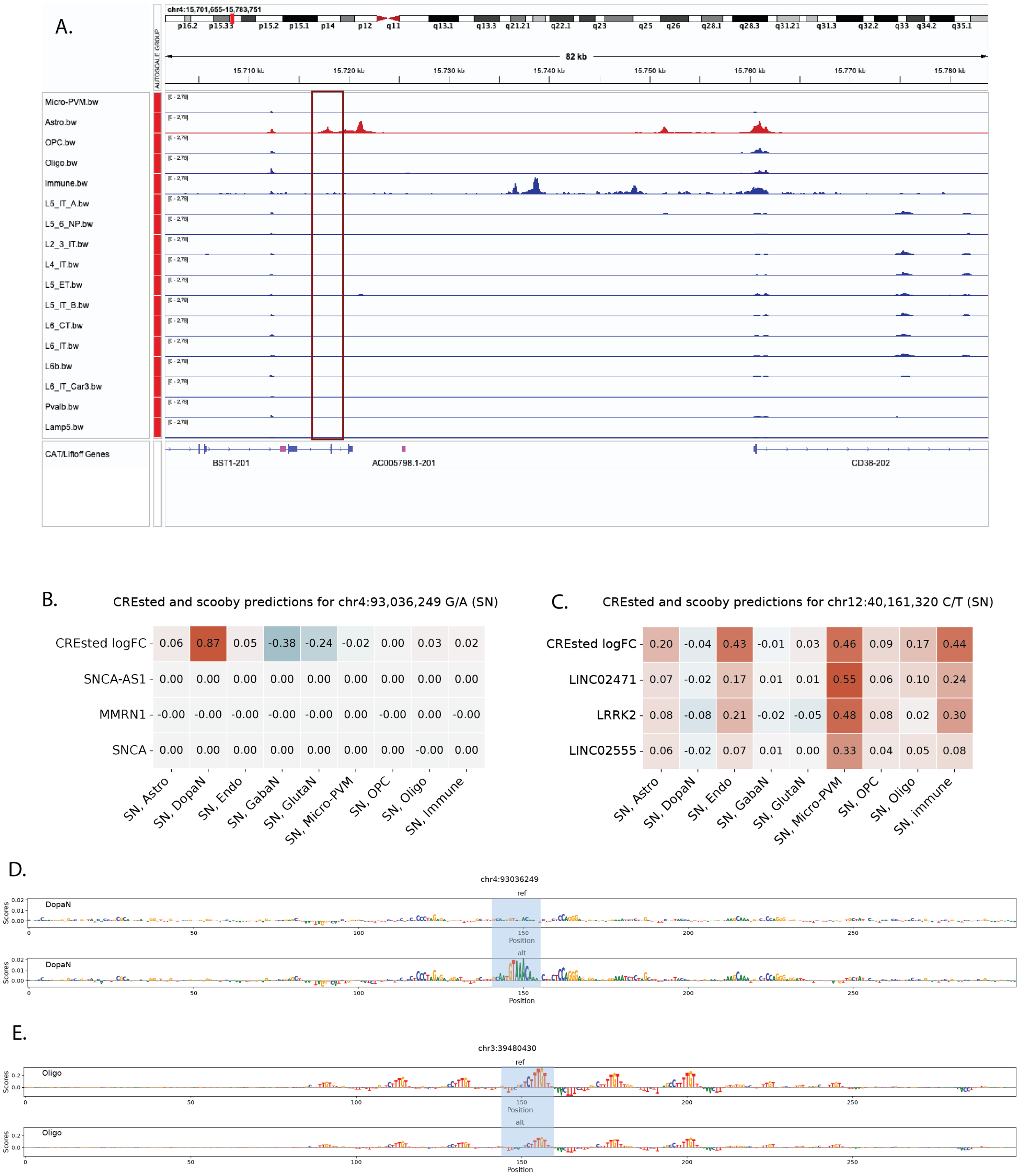


**Supplementary figure 11: Variant-specific quantifications and visualizations**

**A.** IGV view of cell type-specific chromatin accessibility tracks for region chr4:15,701,655-15,783,751. An astrocyte-specific intronic peak in *BST1* is boxed. **B.** CREsted and scooby predictions for variant chr4:93,036,249 G/A in SN cell types. **C.** CREsted and scooby predictions for variant chr12:40,161,320 C/T in SN cell types. **D**. DeepExplainer plots for REF and ALT alleles in Dopaminergic Neurons (SN) for chr4:93,036,249 G/A. **E**. DeepExplainer plots for REF and ALT alleles in Oligodendrocytes (SN) for chr3:39,480,430 G/A.
