## Supplementary tables legend for "Modeling *cis*-regulatory variation in human brain enhancers across a large Parkinson’s Disease cohort"

**Supplementary table1: Donor overview.** Overview of donor metadata.

Columns:
donor_id – internal donor id
age_at_collection – age at which the tissue is collected
sex – sex of the donor
biobank – name of the biobank of origin
PMI – post-mortem interval: time between donor death and tissue collection
PD_status – Parkinson's disease status of the donor. Note: the donor with 'Other neurological disorder' is considered as a control donor in the analysis
brain_region: brain region tissue available, CC: cingulate cortex, SN: substantia nigra, MC: motor cortex
snRNA_technologies – donor transcriptome assays
snATAC_technologies – donor chromatin accessibility assays

**Supplementary table 2: LRS WGS tool versioning.** Reporting versions of the tools used for processing LRS WGS data.

Columns:
Sample – internal region-specific sample ID
bcftools – bcftools version
samtools_cram – samtools version used to create a CRAM file
longphase – longphase version
sniffles2 – sniffles version
clair3 – clair3 version
minimap2 – minimap2 version
samtools_pre_cram – samtools version used for all manipulations except for cram creation
kit – ONT kit used – either SQK-LSK110, or SQK-LSK114
method – Fiber-Seq, or Native (i.e. no modifications introduced, normal LRS WGS protocol)
model_basecalling – ONT model used for basecalling
mod_model_basecalling – what modified bases have been called. If empty, no modified basecalling was performed.
dorado_version – dorado version used. If empty, guppy was used for basecalling.
basecaller – whether guppy or dorado was used for basecalling.
guppy_version – guppy version used. If empty, dorado was used for basecalling.

**Supplementary Table 3: Differential gene expression results.** Reporting results for differential gene expression analysis for all tested cell types. Benjamini-Hochberg p-value correction was applied for each brain region separately.

Columns:
assay – cell type tested
ID – name of tested gene
logFC – log2 fold change
AveExpr – average expression of the gene
T – t-statistic
P.Value – uncorrected p-value
adj.P.Val – Benjamini-Hochberg adjusted p-value
B – B-statistic, log-odds that the gene is differentially expressed
z.std – standardised z-score
region – brain region

**Supplementary Table 4. caQTL, ASCA and CREsted results.** Reporting results for significant caQTL-ASCA variants (53,841 variants, FDR < 0.1 by both tests) in the corresponding cell types. FDR adjusted p-values are reported in caQTL_pval_adj_full and ASCA_pval_adj_full columns (correction performed on full combined set of 10,712,284 variant-cell type combinations). Peak and variant coordinates are provided in CHM13v2 genome version. Rested results are reported for DeepCC and DeepSN models for CC and SN brain regions, respectively

General columns:
variant_id – variant coordinates, REF/ALT allele
variant_type – SNP/indel
peak_id – peak coordinates
brain_region – CC/SN
cell_type, cell_type_formatted – cell type where the variant was tested

tensorQTL-specific columns:
start_distance – Distance to peak start
af — Alternative allele frequency in the study samples.
ma_samples — Number of individuals carrying at least one copy of the minor allele.
ma_count — Total number of minor alleles across individuals.
slope – caQTL slope from tensorQTL
pval_nominal – Nominal p-value from tensorQTL
caQTL_pval_adj_full – FDR adjusted p-value for caQTL

CHT-specific columns (WASP suit, allele-specific part of the test):
LOGLIKE.NULL — Log-likelihood of the CHT under the null hypothesis of no association between test SNP genotype and the target region.
LOGLIKE.ALT — Log-likelihood of the CHT under the alternative hypothesis of association between test SNP genotype and the target region.
CHISQ — Likelihood-ratio test statistic comparing the alternative and null CHT models.
P.VALUE — Nominal p-value for the CHT association test.
ALPHA — Estimated CHT parameter corresponding to the expected contribution of haplotypes carrying the reference allele to read depth in the target region.
BETA — Estimated CHT parameter corresponding to the expected contribution of haplotypes carrying the alternative allele to read depth in the target region.
PHI — Estimated overdispersion parameter for the read-depth component of the CHT model.
TOTAL.AS.READ.COUNT — Total number of allele-specific reads across all individuals included in the test.
REF.AS.READ.COUNT — Total number of allele-specific reads assigned to the reference allele across all individuals included in the test.
ALT.AS.READ.COUNT — Total number of allele-specific reads assigned to the alternative allele across all individuals included in the test.
allelic_imbalance – Calculated as *0.5- ALPHA/(ALPHA + BETA)*
ASCA_pval_adj_full - – FDR adjusted p-value for ASCA

CREsted-specific columns:
diff – Difference between ALT and REF allele predictions
logfc – Log2 ratio between ALT and REF allele predictions

**Supplementary Table 5. meQTL results for caQTL-ASCA peaks.** Reporting meQTL results for significant caQTL-ASCA variants (FDR < 0.1 by both tests) with meQTL FDR < 0.1 (9,469 variants). All peak and variant coordinates are provided in CHM13v2 genome version.

Columns:
variant_id – variant coordinates, REF/ALT allele
peak_id – peak coordinates
start_distance – Distance to peak start
af — Alternative allele frequency in the study samples.
ma_samples — Number of individuals carrying at least one copy of the minor allele.
ma_count — Total number of minor alleles across individuals.
slope – meQTL slope from tensorQTL
pval_nominal – Nominal p-value from tensorQTL
meQTL_pval_adj - FDR adjusted p-values (correction performed on full set of 1,504,086 tested variants).

**Supplementary Table 6. eQTL results for caQTL-ASCA variants.** Reporting eQTL results for significant caQTL-ASCA variants (FDR < 0.1 by both tests) with eQTL FDR < 0.05 (4717 variants, 1121 genes). Peak and variant coordinates are provided in CHM13v2 genome version.

Columns:
variant_id – variant coordinates, REF/ALT allele
gene_id – gene name
peak_id – peak coordinates
cell_type
brain_region
start_distance – Distance to gene start
end_distance - Distance to gene end
af — Alternative allele frequency in the study samples.
ma_samples — Number of individuals carrying at least one copy of the minor allele.
ma_count — Total number of minor alleles across individuals.
slope – eQTL slope from tensorQTL
pval_nominal – Nominal p-value from tensorQTL
eQTL_pval_adj - FDR adjusted p-values (correction performed on full set of 3,395,509,663 tested variant-gene-cell type combinations).

**Supplementary Table 7. Scooby results for caQTL-ASCA variants.** scooby logFC scores for caQTL-ASCA variants. Only variants with absolute scooby_logFC > 0.05 are reported (for the corresponding genes and cell types).

Columns
variant_id – variant coordinates, REF/ALT allele
gene_id – gene name
peak_id – peak coordinates
cell_type
scooby_logfc – logFC from scooby model
brain_region

**Supplementary Table 8. caQTL-ASCA variants in GWAS loci.**
Donor enhancer variants in linkage block boundaries with optional GWAS p-value.

Columns:
variant_id – variant coordinates, REF/ALT allele
lead_variant_rsid – rsID of lead variant of linkage block containing the donor variant
peak_id – T2T/hs1 genomic region of ATAC peak of variant
GWAS_p_val - GWAS p-value of donor variant if variant is also assayed in GWAS

dist_to_lead - distance of donor variant to the lead variant of the linkage block the donor variant is in
Rs_donor - rsID of donor variant
R2 – R2 value between lead and donor variant
GWAS_p_value_lead – GWAS p value of the lead variant
nearest_gene – nearest gene to the lead variant according to GWAS summary

**Supplementary Table 9. Clinvar table with potentially pathogenic variants present in our donor cohort.** We overlapped all donor variants with Clinvar database, as descriped in Methods section ‘Clinvar overlap’.

Columns:
Chr – chromosome
POS – position on the chromosome
ID – clinvar variant ID
REF – reference allele
ALT – reference alternative
INFO – information field from clinvar with various IDs
nHet – number of heterozygous donors
nHomAlt – number of homozygous alternative donors
nHomRef – number of homozygous reference donors
HetSamples – donor IDs for donors who are heterozygous for given variant
HomSamplesAlt – donor IDs for donors who are homozygous alternative for given variant
Link clinvar – link to Clinvar entry of the variant
